# Spatial Patterns of Microbial Diversity, Composition and Community Structure in Fe-Mn Deposits and Associated Sediments in the Atlantic and Pacific Oceans

**DOI:** 10.1101/2022.03.21.485154

**Authors:** Natascha Menezes Bergo, Adriana Torres-Ballesteros, Camila Negrão Signori, Mariana Benites, Luigi Jovane, Bramley J. Murton, Ulisses Nunes da Rocha, Vivian Helena Pellizari

**Affiliations:** Instituto Oceanográfico, Universidade de São Paulo, São Paulo, Brazil; Rothamsted Research, Harpenden, England; National Oceanography Centre, Southampton, England; Helmholtz Centre for Environmental Research, Leipzig, Germany

**Author notes:** **Corresponding author**, Correspondence to Dra Vivian Helena Pellizari. **Availability of data and material**, The sequencing reads generated for this study can be found in the National Centre for Biotechnology Information (NCBI) database under the BioProject PRJNA814217.

**Keywords:** Deep-sea Ferromanganese Crusts and nodules, Microbial Community, Biogeochemical Cycling, Rio Grande Rise, Tropic Seamount, Geomicrobiology

## Abstract

Mining of deep-sea Fe-Mn deposits will remove crusts and nodules in large areas from the seafloor. The growth of a few millimeters of these minerals by Fe and Mn oxides precipitation takes millions of years, and yet little is known about their microbiome. Besides being key elements of the biogeochemical cycles and essential links of food and energy to deep-sea trophic webs, microbes have been identified to affect manganese oxide formation. Hence, polymetallic crusts and nodules may present unique habitats that deserve better understanding. In this study, we determined the composition and diversity of Bacteria and Archaea in deep-sea Fe-Mn crusts, nodules, and associated sediments from two oceanic elevations in the Atlantic Ocean, the Tropic Seamount in the northeast and the Rio Grande Rise (RGR) in the southwest. Sequencing of the 16S rRNA gene was performed using the Illumina MiSeq platform and statistical analyses using environmental data were performed in R. Additionally, we included public domain environmental DNA data of Fe-Mn crusts, nodules, and associated sediments from Clarion-Clipperton Zone and Takuyo-Daigo Seamount in the Pacific Ocean to compare microbial diversity in Fe-Mn deposits from different ocean basins. Our results indicated that Atlantic seamounts harbor an unusual and unknown Fe-Mn deposit microbiome with lower diversity and richness compared to deposits from Pacific areas. Crusts and nodules from Atlantic seamounts revealed the presence of unique taxa (Alteromonadales, Nitrospira, and Magnetospiraceae) and a higher relative abundance of sequences related to potential metal-cycling bacteria, such as Betaproteobacteriales and Pseudomonadales. The microbial beta-diversity from Atlantic seamounts was clearly grouped into microhabitats according to crusts, nodules, and sediments geochemical composition. Furthermore, community structure analysis using principal coordinate analysis also showed that the microbial communities of all seamounts were significantly divided into ocean basins and sampling areas. Despite the time scale of million years for these deposits to grow, a combination of environmental settings (temperature, salinity, depth, substrate geochemistry, nutrient, and organic matter availability) played a significant role in shaping the crusts and nodules microbiome, which was distinct between the Atlantic and Pacific Fe-Mn deposits. Our results suggest that the microbial community inhabiting Fe-Mn deposits participate in biogeochemical reactions indispensable to deep-sea ecosystems, which implies that understanding the microbial community is of utmost importance for any baseline environmental study in areas of potential deep-sea mining.

**Graphical abstract:** 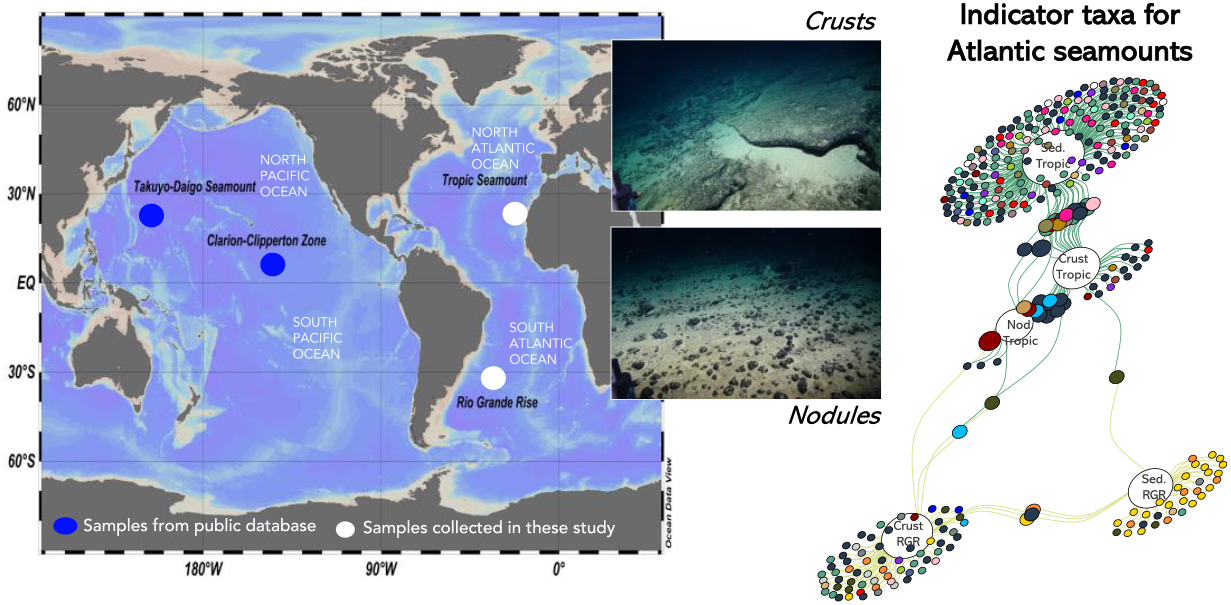

**Highlights:**

- Atlantic deposits showed lower diversity and richness compared to Pacific deposits
- Fe-Mn crusts and nodules are a potential specific ecological niche
- Atlantic Fe-Mn deposits harbor an unusual and unknown microbiome
- Temperature, salinity, depth, and substrate geochemistry at Atlantic Fe-Mn deposits may drive community composition

## INTRODUCTION

Seamounts and oceanic rises support unique biomes in the deep-sea and are globally distributed across the ocean seafloor (Shi *et al*., 2020). They are important habitats for marine species dispersion and evolution (Shi *et al*., 2020). The distinctive oceanographic features of seamounts, such as geographical isolation, topographically induced turbulent water mixing, and rapid current speeds are favorable conditions to establish multiple benthic assemblages (Boehlert & Genin, 1987; Samadi *et al*., 2007). Furthermore, the seamount flanks are frequently covered by volcanic rocks with variable amounts of ferromanganese (Fe-Mn) crusts (Asavin *et al*., 2008; Yeo *et al*., 2018).

Benthic microorganisms are potentially involved in the elemental transition between sea-water and Fe–Mn crusts (Kato *et al*., 2019; Wang & Muller, 2019). Indeed, recently was isolated a chemolithoautotrophic manganese-oxidizing Nitrospirae capable of precipitating Mn (Yu & Leadbetter, 2020). Besides, microorganisms are essential contributors to marine biogeochemical cycles and possibly supply high productivity found in seamounts (Morato *et al*., 2010; Rowden *et al*., 2010; Leitner *et al*., 2020). Bacteria and Archaea have been studied in the deep ocean, where they are diverse and possibly supply the energy flow in the Pacific and Atlantic seamounts (Fullerton *et al*., 2017; Liu *et al*., 2019; Orcutt *et al*., 2020; Bergo *et al*., 2021).

The growing global demand for metals and rare metals has renewed interest in mining Fe-Mn crusts and nodules (Orcutt *et al*., 2020). Deep-sea mining operations will disturb or erode the seafloor and create near-bottom sediment plumes (Miller *et al*., 2018), consequently affecting micro- and macrobenthic life. Despite all the studies that have been conducted since Fe-Mn deposits were discovered in the 1870s (Murray, 1891), little is known about the microbial diversity of seamount environments (Molari *et al*., 2020), especially in the Atlantic Ocean (Bergo *et al*., 2021). Most efforts to study Atlantic seamounts with Fe-Mn deposits have been to assess mineral resources and their megafaunal community (Perez *et al*., 2018; Yeo *et al*., 2018; Montserrat *et al*., 2019; Ramiro-Sánchez *et al*., 2019; Benites *et al*., 2020). Previous studies on the diversity, taxonomy, and functions of microbial communities of seamount with Fe-Mn deposits have mainly focused on the Pacific Ocean (Nitahara *et al*., 2017; Kato *et al*., 2018; Kato *et al*., 2019; Liu *et al*., 2019).

In this study, we investigated the diversity of bacterial and archaeal communities associated with Fe-Mn crusts, nodules, and sediments from the Tropic Seamount (Northeast Atlantic Ocean) and Rio Grande Rise (RGR) (Southwest Atlantic Ocean). We also compared them with other Fe-Mn crusts, nodules, and sediments from the Clarion-Clipperton Zone and Takuyo-Daigo Seamount in the Pacific Ocean. Our main goal was to increase understanding of the (1) microbial diversity in Fe-Mn deposits in seamounts from different ocean basins (Atlantic and Pacific); (2) environmental settings and crust and nodule chemical features that influence the microbial community within Fe-Mn deposits from Atlantic seamounts; (3) differences between community members in Fe-Mn crusts and sediments from Atlantic seamounts; and (4) potential roles of the microbial community in biogeochemical processes of Fe-Mn deposits and associated sediments in Atlantic seamounts. To address these objectives, we collected samples of Fe–Mn crusts, nodules, and sediments from Tropic seamount, as well as crusts and sediments from RGR, and sequenced the 16 rRNA genes on an Illumina platform.

## MATERIALS AND METHODS

### Field Sampling in Atlantic Seamounts

Samples of Fe-Mn crusts, nodules, and associated sediments were collected from the Tropic Seamount (23°55’ N, 20° 45’W) and RGR (31°0’ N, 36° 0’W) in the Northeast and Southwestern Atlantic Ocean, respectively. The Tropic is a flat-topped seamount (guyot) raised from 4000 m to approximately 1000 m water depth and located off the passive continental margin of West Africa (Koschinsky *et al*., 1996; Yeo *et al*., 2018) (Figure 1A). The RGR is an extensive oceanic rise of nearly 150,000 km^2^ rising from approximately 4000 to 600 m water depth, and located about 1,000 km away from the Brazilian coast, between Brazil and Argentine oceanic basins (Cavalcanti *et al*., 2015; Montserrat *et al*., 2019) (Figure 1A).

**Figure 1.**
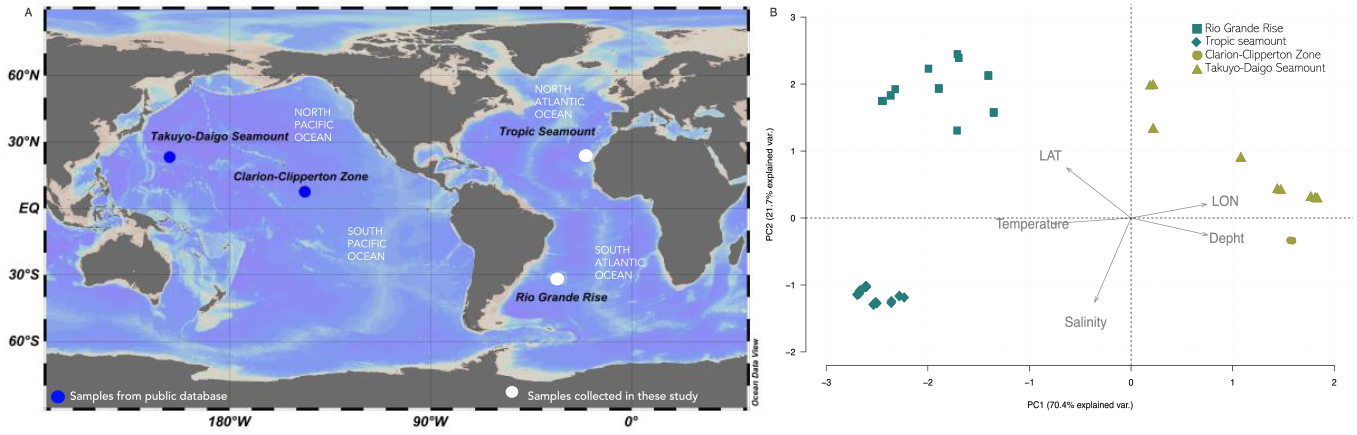
Study areas in the Atlantic and Pacific oceans and principal component analysis for physicochemical parameters. (a) A map of the study regions showing the sampling area from Tropic Seamount in the Northeast Atlantic Ocean and Rio Grande Rise in the Southwest Atlantic Ocean (white circles, collected in this study) and sampling area from Clarion-Clipperton Zone and Takuyo-Daigo Seamount in the North Pacific Ocean (blue circle, downloaded from public databases) (Lindh et al., 2017; Kato et al., 2018). (b) Principal component analysis for samples collected from Rio Grande Rise (square) and Tropic Seamount (diamond) in the Atlantic Ocean (cyan), and Clarion-Clipperton Zone (circle) and Takuyo-Daigo Seamount (triangle) in the Pacific Ocean (pale green). Oceanographic parameters (temperature, salinity, depth, latitude, and longitude) are shown.

Samples were collected in two scientific cruises during the Marine E-Tech project, cruise JC142 onboard the Royal Research Ship James Cook (National Oceanographic Centre, United Kingdom) in October 2016 and cruise RGR1 onboard the Research Vessel Alpha Crucis (Universidade de São Paulo, Brazil) in February 2018 (Supplementary Table 1). On both cruises, Fe-Mn crusts and nodules were aseptically retrieved from the ROV and the dredge area. The surface of each crust and nodule sample was washed with 5 ml of seawater previously filtered with a 0.2 μm-pore polycarbonate membrane to remove loosely attached particles. The sediments were collected by push core and box core, and subsampled into depth intervals of 0-5, 5-10, and 10-15 cm by shipboard subcoring for push core and a sterile spatula in the case of box corer. All samples were stored in sterile Whirl-Pak bags at −80°C.

### Geochemical Analysis of Fe-Mn Crusts, Nodules, and Sediments

We determined the composition of major elements of Fe-Mn crusts, nodules, and associated sediments using a benchtop wavelength dispersive XRF spectrometer Supermini200 Rigaku with a Pd-anode x-ray tube of 50kV and 4mA of power. A scintillation counter was used to detect heavy elements (Ti to U) and a gas-flow proportional counter for light elements (O to Sc). Samples were weighed and heated in an oven at 105°C for 24h to release the sorbed water. Subsequently, samples were powdered using an agate mortar and pestle, sieved through a <150μm mesh, loaded into the XRF equipment and analyzed in a He atmosphere at Centro Oceanográfico de Registros Estratigráficos (CORE), Instituto Ocenográfico, Universidade de São Paulo.

Micronutrients (B, Cu, Fe, Mn, and Zn), organic matter, organic carbon, total nitrogen, nitrate, ammonia, and sulfate were analyzed in the sediment’s samples at Luiz de Queiroz College of Agriculture (Department of Soil Sciences, ESALQ-USP, Brazil) according to methods previously described (Van Raij *et al*., 2001).

### Statistical Analysis of Hydrographic and Geochemical Data

Deepwater hydrographic data (temperature, salinity, and depth) from the Clarion-Clipperton Zone and Takuyo-Daigo seamount were downloaded from the National Center for Biotechnology Information (NCBI) and the DNA Data Bank of Japan (DDBJ) databases, respectively (Lindh *et al*., 2017; Kato *et al*., 2018). We applied principal component analysis (PCA) based on Euclidean distance to the environmental parameters (latitude, longitude, temperature, salinity, and depth) to detect similarities of oceanographic features among oceans. We also used a PCA based on Euclidean distances to determine the influence of different environmental parameters (nitrogen and sulfur compounds, micronutrients, and chemical compositions) to investigate their effect on the sample distributions. For both analyses, the samples were classified according to their sampling location across the oceans. Additionally, we performed the Wilcoxon matched pair using the vegan package to verify differences in oxides, micronutrients, nitrogen, and sulfur compounds concentrations between Tropic Seamount and RGR.

### DNA extraction, 16S rRNA Gene Amplification, and Sequencing

The Fe-Mn crust and nodule surface (0–10 mm) was subsampled using sterile hammers and chisels and then crushed in a sterile agate mortar. DNA was extracted from 5 g of crust, nodule, and sediment aliquots with FastDNA™ SPIN Kit for soil (MPBiomedical), according to the manufacturer’s protocol (Shulse *et al*., 2017; Lindh *et al*., 2017).

Before sending the samples for preparation of Illumina libraries and sequencing, the V3 and V4 region of the 16S rRNA gene was amplified with the primer set 515F (5’ – GTGCCAGCMGCCGCGGTAA - 3’) and 926R (5’ – CCGYCAATTYMTTTRAGTTT - 3’) (Parada *et al*., 2016) to check for the amplification of 16S using the extracted DNA. Negative (no sample) extraction controls were used for PCR amplification and Illumina sequencing to check for the presence of possible environmental contamination (Sheik *et al*., 2018). Illumina DNA libraries and sequencing were performed at MR DNA (Shallowater, TX, United States) on a MiSeq platform in a paired-end read run (2 × 250bp) following the manufacturer’s guidelines. Sequencing outputs were raw sequence data information.

### Sequencing Data Processing and Statistical Analyses

DNA sequence reads from Fe-Mn crust and associated sediments samples from the Pacific Ocean were downloaded from the NCBI SRA database (Lindh *et al*., 2017) and DDBJ SRA database (Kato *et al*., 2018) and processed under the same procedure as the raw sequences generated in this study. The demultiplexed sequences were analyzed with the software package Quantitative Insights Into Microbial Ecology (QIIME 2) version 2019.4 (Bolyen *et al*., 2019). Sequences were denoised using DADA2 (Callahan *et al*., 2016) with the parameters listed in **Supplementary Table 3**. We removed amplicon sequence variants (ASVs) with sequences of less than 10 occurrences. The taxonomy was assigned to the representative sequences of ASVs using a Naive Bayes classifier pre-trained on SILVA release 132 clustered at 99% identity. FastTree and MAFFT (Katoh & Standley, 2013) were used to create a rooted phylogenetic tree used to calculate phylogenetic diversity metrics.

Diversity and phylogenetic analyses were performed with PhyloSeq (McMurdie & Holmes, 2012), ggplot2 (Wickham, 2009), and vegan (Oksanen *et al*., 2013) packages in the R software (Team, 2018). ASVs affiliated with chloroplasts and Eukarya were removed from subsequent analyses. Alpha diversity metrics (e.g., observed sequence variants, Chao1, and Shannon diversity) were calculated based on ASV relative abundances for each ocean and Atlantic seamount. To determine if there were significant differences between alpha diversities, analysis of variance (Kruskal–Wallis one-way ANOVA on ranks test) and subsequent post-hoc Wilcoxon matched pair test were performed in R. ASVs were normalized by variance stabilizing transformation using the R package “DESeq2” (Love *et al*., 2014). Beta diversity among oceans and Atlantic seamounts was analyzed using an ordinated weighted Unifrac normalized distance and visualized using principal coordinate analysis (PCoA, package Phyloseq). We performed PERMANOVA analysis to compare groups in the PCoA plots with the adonis function in the R package vegan, and betadisper was used to assess the differences in dispersions between sample groups (Anderson, 2006). The IndicSpecies identified relative abundance of taxonomic indicators (Cáceres *et al*., 2010). We conducted the analysis was conducted on ASV counts excluding ASVS < 20 reads. We compared the relative abundance and frequency of each ASV to identify those specifically associated with only one substrate (unique) and those whose niche breadth encompasses several substrates (shared). The results were visualized as networks with the igraph R package (Csárdi & Nepusz, 2006).

Distance-based redundancy analysis (dbRDA) was performed to investigate the environmental drivers (depth, temperature, salinity, Al_2_O_3_, Cl, CaO, Co_2_O_3_, Fe_2_O_3_, K_2_O, MgO, MnO, NiO, P_2_O_5_, SiO_2_, SO_3_, TiO_2_, and V_2_O_5_, concentrations) influencing the prokaryotic community structure (package Vegan). Before running the analysis, we normalized the physicochemical characteristics of deep-waters, crusts, nodules and sediments to make the sum of squares equal to one. The ANOVA test was used to test the significance of the dbRDA model and to identify the best set of explanatory variables (p < 0.05) for dbRDA analysis. Raw sequence data generated for this are publicly available in the National Centre for Biotechnology Information (NCBI) database under the BioProject PRJNA814217.

## RESULTS

### Physicochemical Characteristics of Deep-sea Waters, Crusts, Nodules and Sediments

The physicochemical characteristics of deep-sea waters surrounding Pacific Ocean samples were different from those collected in the Atlantic Ocean samples. The temperature ranged from 1.5 to 4°C, and salinity values were between 34.3 and 34.7 psu in deep-sea waters (1432 – 5577 m) surrounding samples from Clarion-Clipperton Zone and Takuyo-Daigo Seamount, (Supplementary Table 1). The temperature was higher in deep-sea waters (685 – 2307 m) overlying the top of the Tropic and RGR seamount than those observed in the Pacific Ocean, ranging from 2.9 to 7.01°C with salinity values between 34.3 and 35.2 psu (Supplementary Table 2). Geochemically, Fe-Mn crusts, and nodules from Tropic Seamount have significantly higher average concentrations of Co_2_O_3_ (1.64 wt%), Fe_2_O_3_ (72.24 wt%), MnO (38.13 wt%), NiO (0.99 wt%), SO_3_ (6.72 wt%), TiO_2_ (0.50 wt%), and V_2_O_5_ (0.15 wt%) when compared to the RGR crusts, (Supplementary Figure 1). Besides that, sediment samples from Tropic Seamount were characterized by significantly higher concentrations of nitrogen and sulfur compounds and micronutrients (NH_4_: 826 mg.kg-1, NO_3_: 82.50 mg.kg-1, N_total_: 861 mg.kg^−1^, SO_4_^−2^: 992 mg.kg^−1^, B: 3.74 mg.kg^−1^, Cu: 0.50 mg.kg^−1^, Fe: 4.50 mg.kg^−1^, Mn: 2.75 mg.kg^−1^, and Zn: 0.40 mg.kg^−1^) when compared to the RGR sediments (Supplementary Figure 2).

Spatial variations between Pacific and Atlantic Ocean samples were well explained (92.1%) by environmental characteristics (longitude, latitude, temperature, and depth) in the PCA analysis, (Figure 1B and Supplementary Table 4). Samples were clustered into two categories: oceans and sampling areas within each ocean basin (Figure 1B). Spatial variations between Atlantic Ocean samples were well explained (75.6%) by environmental characteristics (longitude, latitude, salinity and CaO, Fe_2_O_3_, MnO, Al_2_O_3_, TiO_2_, V_2_O_5_, Co_2_O_3_, and NiO) in the PCA analysis (Supplementary Figure 3A and Supplementary Table 5). Samples were clustered into two categories: the two sampling areas in the Atlantic Ocean and the type of substrate (Supplementary Figure 3A). In addition, spatial variations between Atlantic Ocean sediment samples were well explained (69.1%) by nitrogen and sulfur compounds (N_total_, NH^4+^, NO^−3^, and SO_4_^−2^), and micronutrients (B, Mn, and Zn) in the PCA analysis (Supplementary Figure 3B and Supplementary Table 6).

### Alpha and Beta Diversity Estimates in the Pacific and Atlantic Oceans

We obtained 10,573,651 DNA sequences from 113 samples, 3,969,611 for the Atlantic Ocean and 6,604,040 for the Pacific Ocean. After filtering low-prevalence features and eukaryotes, we obtained 19,970 amplicon sequence variants, 5,783 for the Atlantic Ocean and 14,187 for the Pacific Ocean. Microbial alpha diversity indices were significantly higher in samples from the Pacific Ocean (Kruskal-Wallis test, p < 0.01; Figure 2A and Supplementary Table 7) than samples from the Atlantic Ocean. Pairwise comparisons of Shannon and Chao1 indices indicated the alpha diversity of sediments and nodules from the Pacific Ocean were significantly higher (Wilcoxon test, p < 0.01; Figure 2A); and (ii) the Chao1 index of crusts did not significantly differ between oceans (Wilcoxon test, p > 0.05; Figure 3). Shannon and Chao1 indices were significantly higher in sediment samples from Tropic seamount when compared with RGR (Kruskal-Wallis test, p < 0.01; Supplementary Figure 4A and Supplementary Table 8). Pairwise comparisons of the microbial communities inhabiting the same substrate indicated the alpha diversity in 0-5 and 5-10 cm sediment layers were significantly higher compared to other samples (Wilcoxon test, p < 0.01; Supplementary Figure 4A). In addition, the alpha diversity in crusts and 10-15 cm sediment layer did not significantly differ between Atlantic seamounts (Wilcoxon test, > 0.05; Figure 2A).

**Figure 2.**
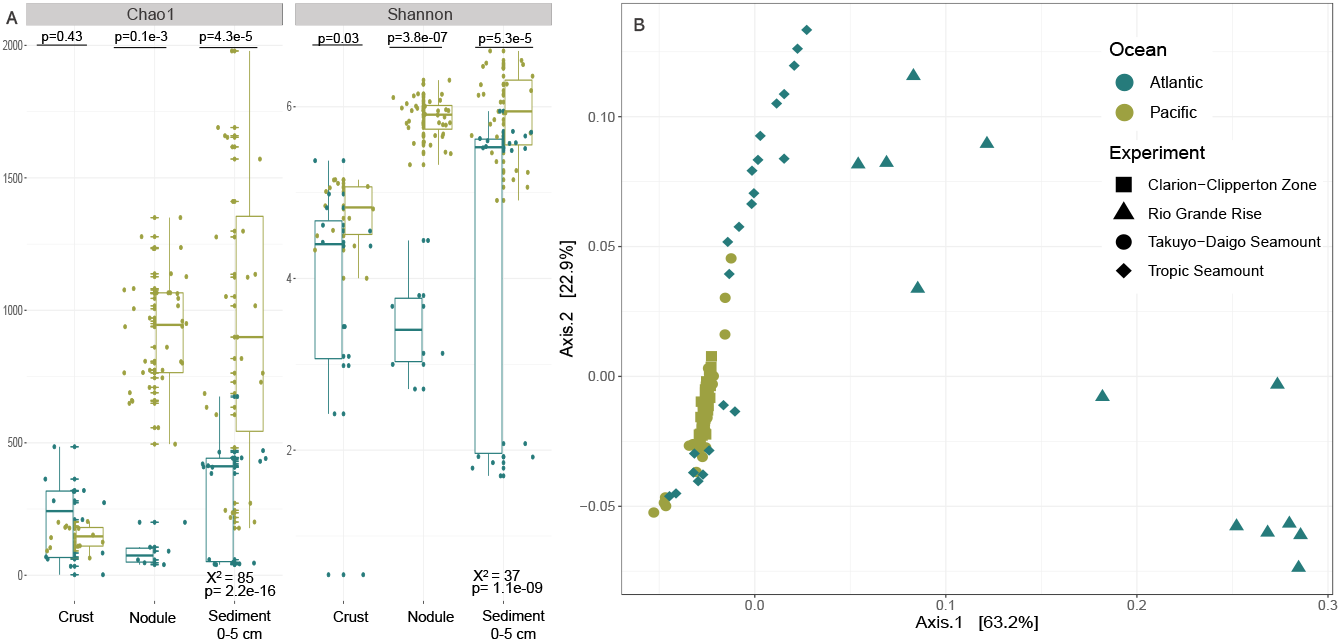
Alpha and beta diversity of microbial communities in the Fe-Mn crusts, nodules, and associated sediment from Atlantic (cyan) and Pacific (pale green) Ocean. (a) Alpha diversity (Chao1 and Shannon indexes) medians of microbial communities. Means were compared by the Kruskal-Wallis test. Pairwise comparisons were performed using the Wilcoxon test. (b) Beta diversity principal coordinate analysis (PCoA) based on the ordinated weighted Unifrac normalized distance. Fe-Mn crust, nodule, and associated sediment from RGR (triangle) and Tropic Seamount (diamond).

**Figure 3.**
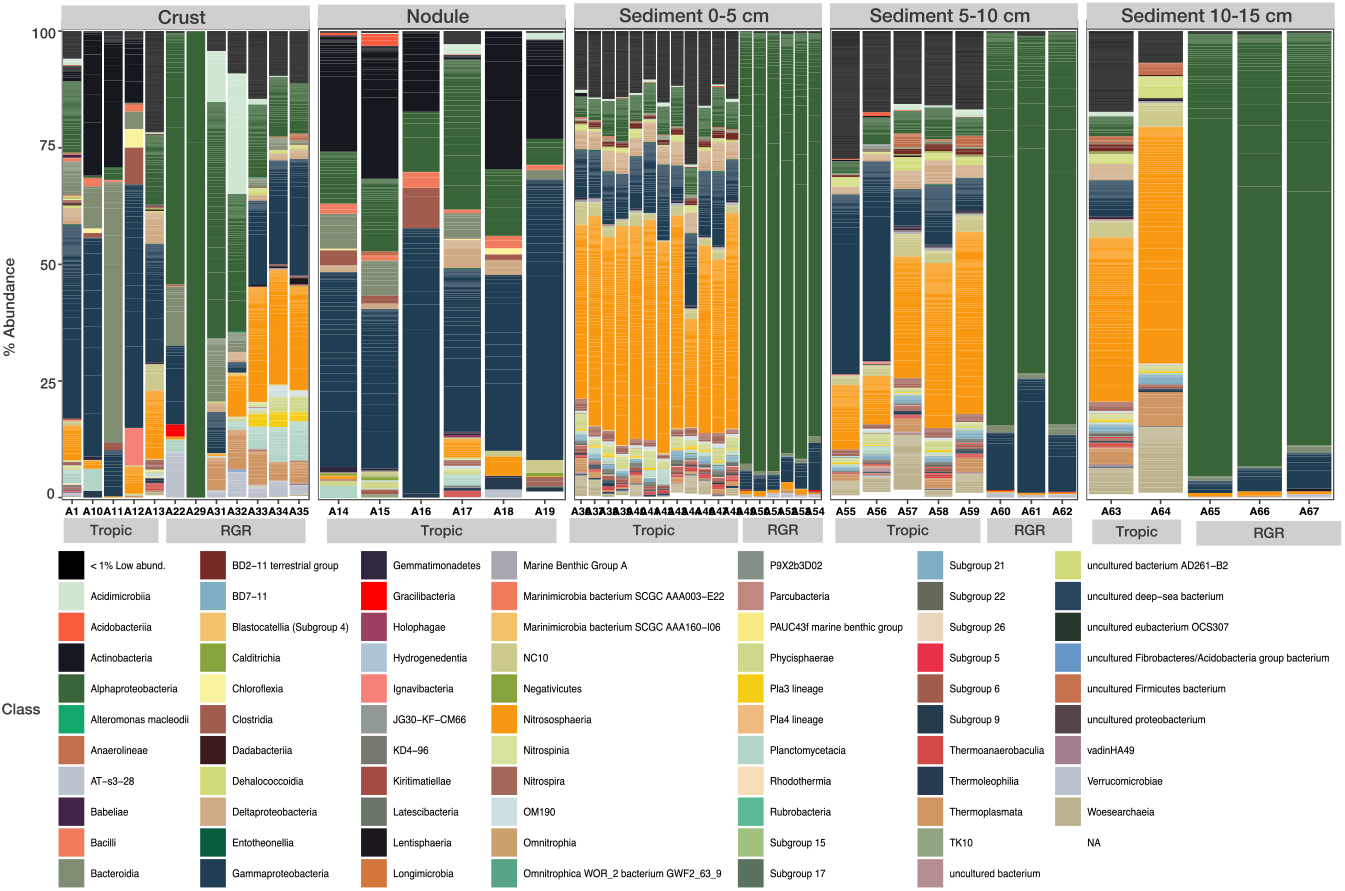
Relative abundances of bacterial and archaeal taxonomic composition for class in the Fe-Mn crusts, nodules, and sediment depth intervals 0-5 cm, 5-10 cm, and 10-15 cm from Tropic seamount and Rio Grande Rise. Only classes with more than 0.1% abundance are represented. Classes with relative abundances below 5% were grouped for the low abundance groups. Gray boxes at the top indicate sample substrates and gray boxes at the bottom differentiate samples from Tropic seamount (Tropic) and Rio Grande Rise (RGR).

Microbial beta-diversity explored by ordinated weighted Unifrac normalized distance did not reveal a clear distinction between Pacific and Atlantic samples (Figure 2B). The PCoA analysis captured 86.1% of the total variation of the prokaryotic community composition in the investigated samples. However, analysis of variance with PERMANOVA showed that samples differed significantly when comparing Pacific and Atlantic Oceans (adonis, df=1, F=13.8, r^2^ = 0.09, p = 0.001) and type of substrate (adonis, df=2, F=4.6, r^2^ = 0.06, p = 0.001). The BETADISPER homogeneity test among the groups showed a significant (p < 0.05) difference in group dispersions for a different type of substrate (betadisper, df=2, F=9.9, p = 0.0001), but not for different geographic location (betadisper, df=1, F=0.2, p = 0.6). PERMANOVA and betadisper significant variation indicate that samples under each factor represent a significant difference in composition and homogeneity. Besides, there was a higher similarity between samples grouped according to their geographic location (one-way ANOSIM; R = 0.56, p = 0.001).

In the case of Atlantic seamounts, beta-diversity explored by ordinated weighted Unifrac normalized distance revealed a clear distinction between the studied regions (Supplementary Figure 4B). The PCoA analysis captured 88.3% of the total variation of the community composition. Significant difference was found comparing Tropic and RGR geographic location by both the PERMANOVA (adonis, df=1, F=10.3, r^2^ = 0.20, p = 0.001) and the BETADISPER (betadisper, df=1, F=6, p = 0.02), as well as when comparing the type of substrate (PERMANOVA: adonis, df=2, F=4, r^2^ = 0.16, p = 0.001; betadisper, df=2, F=3, p = 0.03). The results yielded in the PERMANOVA and the betadisper analyses indicate that selected classes exhibit different groups concerning their composition and homogeneity. In addition, there was a higher similarity between samples grouped according to their type of substrate (one-way ANOSIM; R = 0.28, p = 0.001).

### Patterns in Microbial Community Composition at Atlantic Seamounts

Microbial communities in samples from Tropic seamount and RGR were overall dominated by the phyla Proteobacteria (classes Gamma- and Alphaproteobacteria), Thaumarchaeota, Actinobacteria, and Bacteroidetes. These phyla accounted for a cumulatively average of 83% of the sequences within each sample (Supplementary Figure 5). For example, RGR crusts were dominated by Alphaproteobacteria (50-12%, orders SAR11 clade and Rhodospirillales), Gammaproteobacteria (27-3%, orders Betaproteobacteriales, Pseudomonadales and Alteromonadales), and Nitrososphaeria (26-9%, order Nitrosopumilales) (Figures 3 and Supplementary Figure 7). On the other hand, crusts and nodules samples from Tropic Seamount showed similar group compositions with a dominance of Gammaproteobacteria (46-31%, orders Betaproteobacteriales, MBMPE27, and Pseudomonadales) followed by Actinobacteria (19-16%, order Propionibacteriales), Bacteroidia (22-4%, order Flavobacteriales), and Alphaproteobacteria (15-7%, orders Rhodovibrionales and Rhizobiales) (Figures 3 and Supplementary Figure 6).

Prevalent taxa in sediments from Tropic and RGR was structured differently (Figure 3 and Supplementary Figure 6). A higher number of classes were detected in the sediment layers from Tropic when compared to sediment from RGR. Classes that were more abundant in the sediment layers from RGR included: Alphaproteobacteria (order SAR11 clade), Gammaproteobacteria (order Alteromonadales), and Nitrososphaeria (order Nitrosopumilales), (Figures 3 and Supplementary Figure 6). The microbial community from the sediment layers from Tropic was mostly dominated by Nitrososphaeria (order Nitrosopumilales) flowed by Gammaproteobacteria (orders Betaproteobacteriales, Pseudomonadales, MBMPE27, and Steroidobacterales), Alphaproteobacteria (order Rhodovibrionales), Deltaproteobacteria (order NB1-j), NC10 (order Methylomirabilales), Dehalococcoidia (SAR202 clade), Subgroup 21, and unclassified Woesearchaeia (Figures 3 and Supplementary Figure 6).

The unique ASVs in sediments from RGR mainly belonged to the uncultured Rickettsiales order (n=22), SAR11 clade (n=6), unclassified Alteromonadales (n=2), and Nitrosopumilales (n=2), while in crusts from RGR, the unique ASVs belonged to the Nitrosopumilales order (n=11), unclassified Marine Group II order in Thermoplasmata (n=6), SAR11 clade (n=5), Alteromonadales (n=5), uncultured Bdellovibrionales (n=4), and SAR202 clade (n=2) (Figure 4B). In the Tropic sediments the unique ASVs belonged mostly to the orders Nitrosopumilales (n=45), uncultured Rhodovibrionales (n=14), unclassified Alphaproteobacteria (n=12), NB1-j (n=11), uncultured Nitrospirales (n=11), Steroidobacterales (n=8), Phycisphaerales (n=7), uncultured Methylomirabilales (n=5), uncultured Cytophagales (n=5), Nitrosococcales (n=5), uncultured Subgroup 6 (n=5), unclassified Woesearchaeia (n=5), uncultured Omnitrophicaeota (n=4), uncultured Thermoplasmata (n=4), Betaproteobacteriales (n=4), uncultured BD2-11 terrestrial group (n=4), unclassified EPR3968-O8a-Bc78 (n=3), uncultured Subgroup 21 (n=3), and the uncultured AT-s2-59 (n=3). In contrast, crusts and nodules from Tropic harbored fewer unique ASVs (n=22 and n=3, respectively), most associated to Betaproteobacteriales, Nitrospirales, uncultured Rhodospirillales, SAR324, and the uncultured PAUC43f marine benthic group (within the group ‘Others’) (Figure 4B).

**Figure 4.**
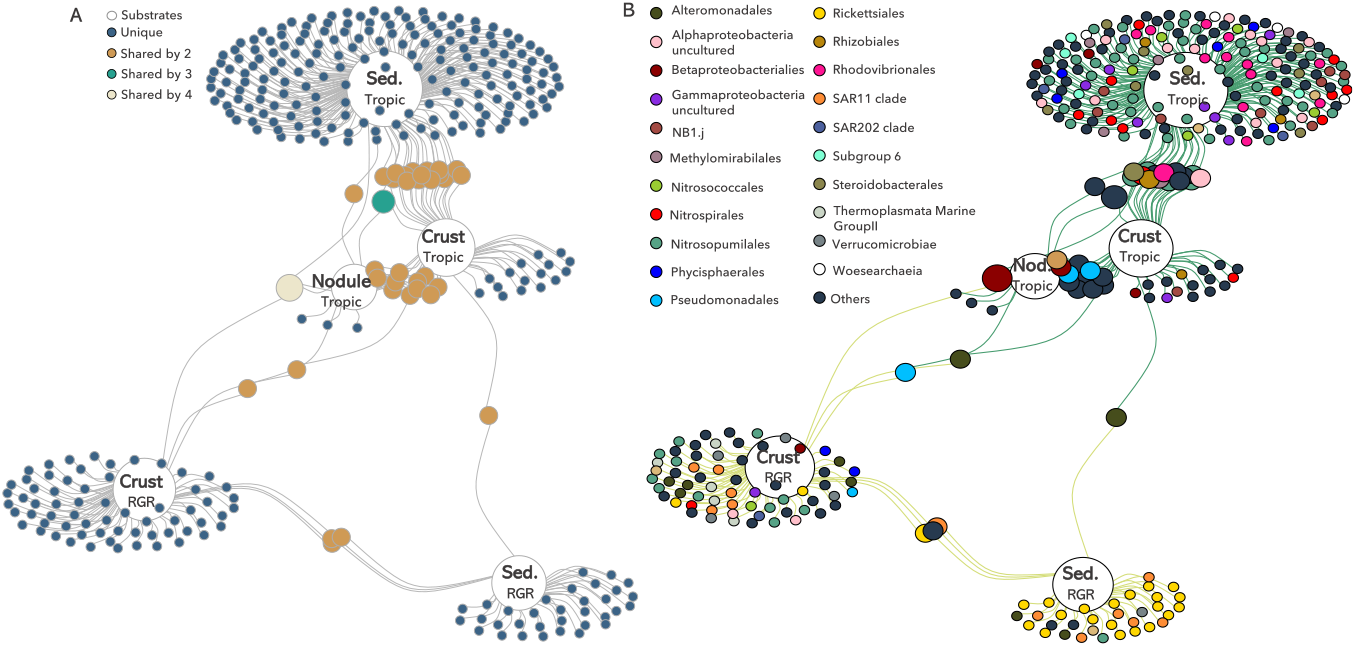
Indicator Species analysis of Fe-Mn crusts, nodules, and sediment (sed) from Tropic seamount (Tropic) and Rio Grande Rise (RGR). The networks (A) indicate the correlation (represented by edges) of unique (1 substrate) and shared ASVs (2 to 4 different substrates); (B) highlight the taxonomic classification of the nodes that belonged to the most represented orders (> 1% of relative abundance in at least one sample). Non-dominant taxa (< 1%) are reported as “Others”. The nodes size reflects “fidelity” to the substrates so that a large node indicates ASVs always present in that substrate.

All substrates from Tropic shared high numbers of ASVs: crusts shared 20 ASVs with sediment (mostly Dadabacteriales, Methylomirabilales Nitrospirales, Nitrosopumilales, Rhodospirillales, and Steroidobacterales), and 11 ASVs with nodules (Pseudomonadales, Micrococcales, Actinomycetales, within the group ‘Others’) (Figure 4). Sediments and crusts from RGR and sediments and nodules from Tropic shared fewer ASVs (n=4 and n=1, respectively), belonging to the Rickettsiales order, SAR11 clade, Flavobacteriales and Thiotrichales orders (within the group ‘Others’), and the MBMPE27 order, respectively (Figure 4). The two seamounts shared fewer ASVs between the substrates: sediment from RGR and crust from Tropic (n=1, Alteromonadales order), and crusts from RGR and Tropic (n=1, Alteromonadales order) (Figure 4). All substrates from Tropic shared 2 ASVs, belonging to the MBMPE27 (within the group ‘Others’) and Rhizobiales orders. All substrates from Tropic shared 1 ASV with RGR crusts (Betaproteobacteriales order).

#### Environmental Drivers of the Microbial Community Structure among Atlantic Seamounts

We investigated the role of the oceanographic parameters shaping microbial community composition using distance-based redundancy analyses, constrained with significant variables, (Figure 5). This analysis showed that oceanographic parameters (depth, temperature, salinity, Al_2_O_3_, Cl, CaO, Co_2_O_3_, Fe_2_O_3_, K_2_O, MgO, MnO, NiO, P_2_O_5_, SiO_2_, SO_3_, TiO_2_, and V_2_O_5_, concentrations) divided the community structure into two clusters (geographic location and type of substrate), and together explained 66.8% of the total variance. The first axis explained most of the variation, which accounted for 32.5% of the variation and was divided into seamounts, i.e., Tropic and RGR. The second axis explained 19.1% of the variation and divided into crust, nodule and sediment samples. Temperature, salinity, depth, CaO, and SiO_2_ were the set of environmental variables that best explained the variations in the community’s structure (p <0.001) (Figure 5). The ANOVA test revealed that Cl was the main environmental factor affecting the distribution of microbes in the sediment (p<0.05) while temperature and salinity, V2O, MnO, and Fe2O3 were the factors affecting crusts and nodules microbial distribution (p<0.05) on Tropic seamount (Figure 5). In samples from RGR, the ANOVA test revealed that depth and CaO were the main environmental factor affecting sediment microbial distribution. At the same time, while concentration of SiO_2_ was factors affecting crust microbial distribution (p=0.009 and p=0.001, respectively, Figure 5). The outcome of this analysis confirmed and strengthened the ANOSIM results.

**Figure 5.**
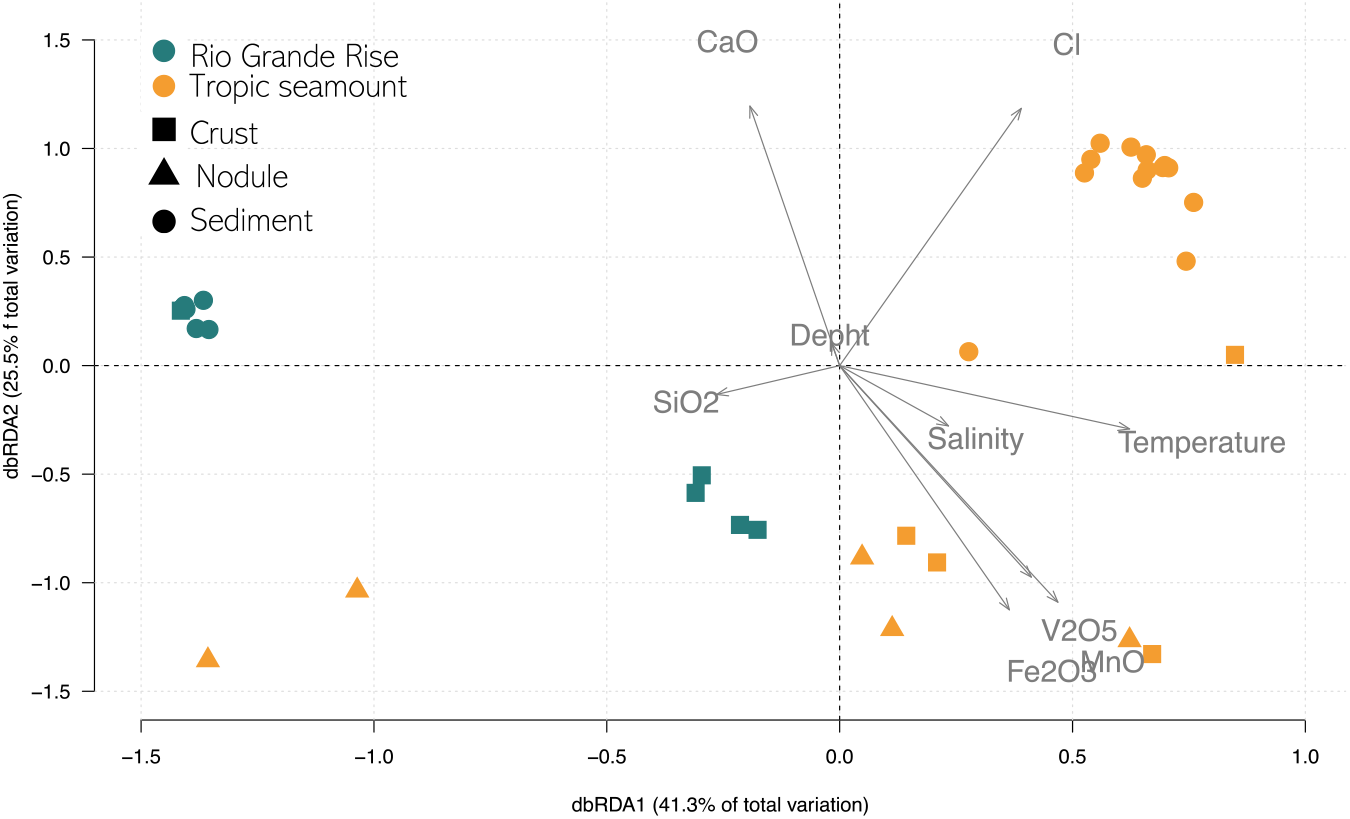
Relationships between environmental variables and prokaryotic community composition from Tropic seamount (orange) and Rio Grande Rise (cyan). The dbRDA axes explain a cumulative 66.8% of the variance in microbial community structure.

## DISCUSSION

### Microbial Composition of Atlantic Seamounts Compared with Other Deep-sea Fe-Mn deposits

Benthic bacterial assemblages in the crust, nodules, and sediments from the Atlantic seamounts showed typical dominance of the classes Gammaproteobacteria, Alphaproteobacteria, Actinobacteria, Bacteroidia, and Deltaproteobacteria, as previously reported for the Pacific Fe-Mn deposits (Wu *et al*., 2013; Lindh *et al*., 2017; Shulse *et al*., 2017). However, we detected significant differences in the microbial community composition between the Tropic seamount and RGR and Pacific Fe-Mn deposits at a lower taxonomic level. For example, the higher relative abundance of the orders Alteromonadales, BD2-11 terrestrial group, Betaproteobacteriales, NC10, MBMPE27, Rhodovibrionales, SAR11, and Steroidobacterales, and the lower relative abundance of Flavobacteria and Bacilli in the Atlantic seamounts differed from the microbial community composition observed at CCZ and Takuyo-Daigo seamount (Lindh *et al*., 2017; Kato *et al*., 2018).

Previous studies have described that Mn-reducing bacteria, such as *Shewanella* and *Colwellia*, are key microorganisms acting to dissolve Mn in Fe–Mn crusts (Blöthe *et al*., 2015). Although we have not identified these bacterial species, the indicator taxa from crusts and nodules detected groups potentially related to metal cycles, such as uncultured Magnetospiraceae (order Rhodospirillales) and Nitrospira (Nitrospirales order). Members of the Nitrospira phylum use Mn as an energy source to grow and produce small nodules of Mn oxide (Yu & Leadbetter, 2020). Besides, members of the family Magnetospiraceae are capable of magnetotaxis and iron reduction (Matsunaga *et al*., 1991). Further, we detected a relatively high abundance of ASVs in the Fe-Mn crusts and nodules associated with Fe reducers and Mn oxidizers from the Alteromonadaceae, Burkholderiaceae, and Pseudomonadaceae families. Representatives of these groups have been proposed as potential keys in forming and growing Fe-Mn crusts and nodules in the Pacific Ocean (Wu *et al*., 2013; Blöthe *et al*., 2015; Kato *et al*., 2018; Hassan *et al*., 2020).

In addition, the indicator taxa result for crusts and nodules identified groups that were normally associated with reduced inputs of organic matter, specifically SAR202 and SAR324. Representatives of the heterotrophic SAR202 and chemolithoautotrophic SAR324 have been proposed as potential indicators of deep water oligotrophic conditions (Galand *et al*., 2010; Orcutt *et al*., 2011; Landry *et al*., 2017). These SAR groups are related to the deep carbon and sulfur cycles, and the SAR202 has been suggested as an important consumer of deep ocean recalcitrant dissolved organic matter (Wei *et al*., 2020). SAR202 and SAR324 were previously described as benthic microbial assemblages in other Fe-Mn deposits (Wu *et al*., 2013; Blöthe *et al*., 2015; Walsh *et al*., 2016; Lindh *et al*., 2017; Molari *et al*., 2020).

Unexpectedly, the SAR11 clade was detected only in the RGR seamount. Closely related sequences of SAR11 were reported previously in sediments across the São Paulo Plateau (Queiroz *et al*., 2020) and crusts from RGR (Bergo *et al*., 2021). SAR11 includes carbon-oxidizing bacteria with multiple depth-specific ecotypes (Cameron *et al*., 2014; Giovannoni *et al*., 2017). Two factors may explain the higher abundance of SAR11 clade I in RGR, the microbial transport through the water column (Hamdan *et al*., 2013; Walsh *et al*., 2016) and/or the influence of surrounding seawater during the crusts and sediments sampling.

Recently, we demonstrated a large proportion of unknown species and hidden functions in the RGR seafloor microbiome (Bergo *et al*., 2021). In fact, the indicator taxa for sediments in both RGR and Tropic seamount reveals an unknown deep-sea environment influenced by unclassified and uncultured groups. Many of these taxa include the unclassified Gammaproteobacteria (order MBMPE27), BD2-11 terrestrial group (phylum Gemmatimonadetes), Rickettsiales, Subgroup 21 (phylum Acidobacteria), and the genus *Woesia* (family Woeseiaceae). There are no cultivates for MBMPE27 and BD2-11 terrestrial groups, and their function remains unknown. However, sequences of these unclassified and uncultured groups have been described at Pacific Fe-Mn deposits (Liao *et al*., 2011; Molari *et al*., 2020; Wu *et al*., 2013).

Moreover, members of the Nitrososphaeria family represented a small portion of total sequences (approximately 0.8%) retrieved from RGR compared to Tropic seamount (approximately 37%). In addition, the indicator taxa result identified a higher number of unique ASVs belonging to Nitrosopumilales order in sediments from Tropic seamount. A high proportion of these potentially chemolithoautotrophic archaea was previously reported for Takuyo-Daigo deep-sea regions (Kato *et al*., 2018). Some authors consider the presence of Nitrososphaeria in energy-limited deep-sea environments as a food web supported by ammonia oxidation and carbon fixation (Tully & Heidelberg, 2013; Zhang *et al*., 2015). Overall, these findings suggest that bacterial and archaeal groups adapted to lithic substrates preferentially colonize crusts and nodules, likely favored by manganese and iron availability.

### Factors Influencing Patterns of Microbial Community Composition and Structure

Studies have shown that microbial community structure can be relatively similar in analogous marine environments even though distancing thousands of kilometers away (Agogué *et al*., 2011; Walsh *et al*., 2015). Nevertheless, the microbial community can also be variable over only a few tens of kilometers away within heterogeneous environments (Hewson *et al*., 2007; Kato *et al*., 2018; Liu *et al*., 2019). In this study, PCA analysis showed that Fe-Mn crusts, nodules, and associated sediment were highly divided into oceans and sampling areas depending on oceanographic parameters from surrounding deep-sea waters. Moreover, the samples from the Atlantic seamounts were highly clustered into crust, nodule, and sediment geochemical composition. These findings indicate environmental homogeneity between the Fe-Mn deposits from Atlantic and Pacific seamounts but significant heterogeneity between Fe-Mn crusts, nodules, and sediments from seamounts in the same ocean (Atlantic). Per diversity analysis using PCoA, the microbial community structure of Atlantic seamounts showed a similar partitioning, supporting an association between the microbial community and local oceanographic parameters.

Our results fell in line with previous observations that higher microbial richness and diversity in marine sediments samples is higher than in Fe-Mn crusts and nodules (Lindh *et al*., 2017; Shulse *et al*., 2017; Kato *et al*., 2018). The higher diversity found in marine sediment samples may occur because of the lower availability of energy sources (e.g., organic matter) in crusts and nodules when compared to sediments (Tully & Heidelberg, 2013). Equally important, the nodules and crusts offer hard substrates and metals, which can select specific microorganisms (Morali *et al*., 2020). In addition, our samples from Atlantic seamounts displayed significantly lower microbial alpha diversity when compared to Pacific areas, indicating that Takuyo-Daigo seamount and CCZ have a different microbial community from those in the Tropic seamount and RGR. However, Shannon diversity for crusts and nodules collected from Tropic seamount and RGR fall within the range of that observed for crusts and nodules from Takuyo-Daigo seamount and CCZ.

Two factors may be shaping the observed community structure and composition variation in Atlantic Fe–Mn crusts. The first is the geochemistry of the different crusts, such as the higher concentrations of cobalt, iron, manganese, nickel, and sulfur in the Tropic crusts. In contrast, crusts from RGR crusts are relatively enriched in aluminum, potassium, and magnesium. Besides geochemical characteristics, microbial community composition and structure patterns on seamounts may also be influenced by oceanographic parameters (water depth, oxygen, and location). This has been previously described for Fe-Mn deposits from the Pacific Ocean and RGR (Nitahara *et al*., 2017; Kato *et al*., 2018; Liu *et al*., 2019; Benites *et al*., 2020; Morali *et al*., 2020).

#### Final Considerations

Our results revealed that: (1) the prokaryotic diversity and community structure in the Atlantic seamounts differed somewhat from the Pacific seamounts, suggesting that benthic microbes reflect regional differences in hydrography and biogeochemistry; (2) the vast array of uncultured and unclassified microbes in this study confirmed the lack of knowledge regarding the deep seafloor microbiome and its link with the biogeochemical processes in the dark Atlantic Ocean; (3) alpha and beta diversity at crust and nodule from the Atlantic seamounts differed from sediment samples, suggesting that Fe-Mn crusts and nodules are a potential specific ecological niche; and (4) the microbial community in the Atlantic seamounts may be sustained by dissolved nitrogen compounds (ammonia) and by sinking particles containing organic carbon and sulfur as energy sources, which are supplied by the photosynthetic ecosystem at the ocean surface. Overall, these insights provided by the prokaryotic community structure emphasize the value of incorporating microbiological surveys more broadly into field sampling campaigns in the Atlantic Fe-Mn deposits. This study establishes a baseline for Atlantic Fe-Mn benthic microbial life that will be critical to support further spatial and/or temporal studies.

Future research will improve our understanding of how the biological communities may respond to disturbances produced by deep-sea mining activities and other possible impacts (as climate change).

## Acknowledgments

We thank the captain and crew of the Royal Research Ship James Cook (NERC, UK) and Research Vessel Alpha Crucis (USP, BR), and scientists who joined the expeditions Marine E-Tech JC142 and RGR1 for data and sampling support. Also, we thank Pedro M. Tura and Carolina L. Viscarra for assistance with sampling and scientific support onboard and all MarineE-tech members for collaboration. Also, we thank Felipe Borim Correa and Rodolfo Toscan for scientific support in processing all environmental DNA data. We thank Linda Waters for the English language review. We thank Rosa C. Gamba, the LECOM research team (IOUSP, São Paulo, Brazil), and the Microbial Data Science research team (UFZ, Leipzig, Germany) for their scientific support.

## Formatting of funding sources

This work was supported by the São Paulo Research Foundation (FAPESP), grant number 14/50820-7, project “Marine ferromanganese deposits: a major resource of E-Tech elements”, which is an international collaboration between the Natural Environment Research Council (NERC, UK) and FAPESP (BRA). NMB was financed by a Doctorate’s fellowship from the Coordenação de Aperfeiçoamento de Pessoal de Nível Superior - Brasil (CAPES) – Finance Code 001 and by the National Council for Scientific and Technological Development – CNPq (grant number 203915/2018-6). CNS was supported by a Postdoctoral fellowship from FAPESP (Grant number 2016/16183-5).

## CRediT authorship contribution statement

Natascha Menezes Bergo: Conceptualization, Methodology, Investigation, Data curation, Writing – original draft, Writing – review & editing.

Adriana Torres-Ballesteros: Formal analysis, Data curation, Writing – original draft, Writing – review & editing.

Camila Negrão Signori: Conceptualization, Writing – original draft, Writing – review & editing, Visualization.

Mariana Benites: Formal analysis, Data curation, Writing – review & editing.

Luigi Jovane: Methodology, Writing – original draft, Writing – review & editing.

Bramley J. Murton: Methodology, Writing – review & editing, Project administration, Funding acquisition.

Ulisses Nunes da Rocha: Conceptualization, Methodology, Writing – review & editing, Visualization.

Vivian Helena Pellizari: Conceptualization, Methodology, Writing – review & editing, Supervision.

## SUPPLEMENTARY MATERIAL

### Supplementary Text

During the JC142 cruise, the remotely operated vehicle (ROV) Isis (National Oceanographic Center, UK) collected Fe-Mn crusts, nodules, and associated sediment samples and measured temperature and salinity using a CTD sensor during sampling dives. Fe-Mn crust and nodule samples were collected using the ROV manipulator arm, while the associated sediment was sampled using a push-core sediment sampler. On the RGR1 scientific cruise, Fe-Mn crusts and sediment samples were collected by dredge and box corer, respectively. Further details about the RGR1 scientific cruise are described by Jovane et al. (2019).

Negative (no sample) DNA extraction controls were used to ensure the extraction quality. DNA integrity was determined after electrophoresis in 1% (v/v) agarose gel prepared with TAE 1X (Tris 0,04M, glacial acetic acid 1M, EDTA 50mM, pH 8), and staining with Sybr Green (Thermo Fisher Scientific, São Paulo, Brazil). DNA concentration was determined using the Qubit dsDNA HS assay kit (Thermo Fisher Scientific, São Paulo, Brazil) following the manufacturer’s instructions.

**Supplementary Figure 1.**
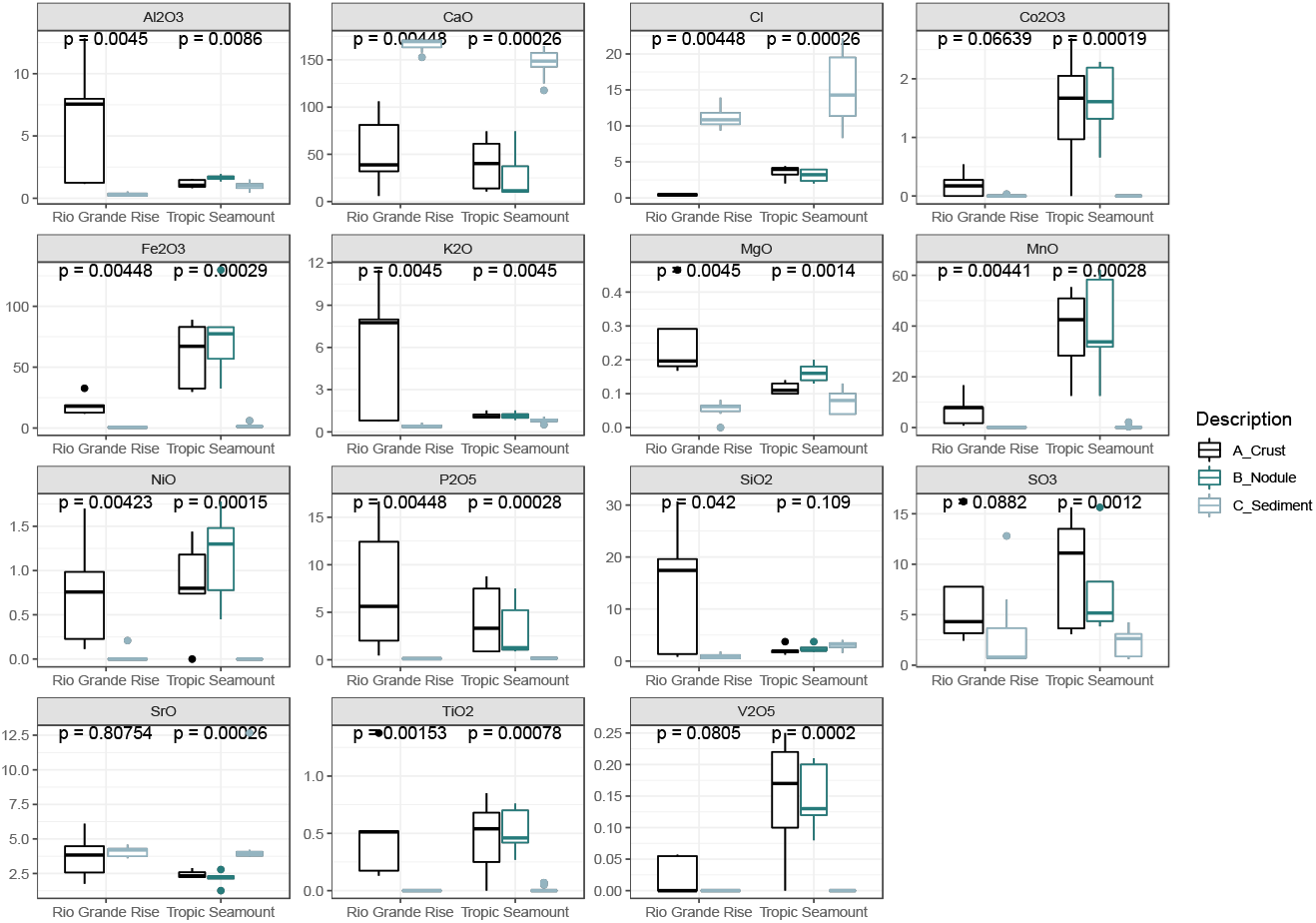
Boxplot of compounds of concentrations in Fe-Mn crust (black), nodule (green), and sediment (gray) from Tropic Seamount and Rio Grande Rise. Note that the vertical scale bars differ between graphs. Pairwise comparisons were performed using the Wilcoxon test.

**Supplementary Figure 2.**
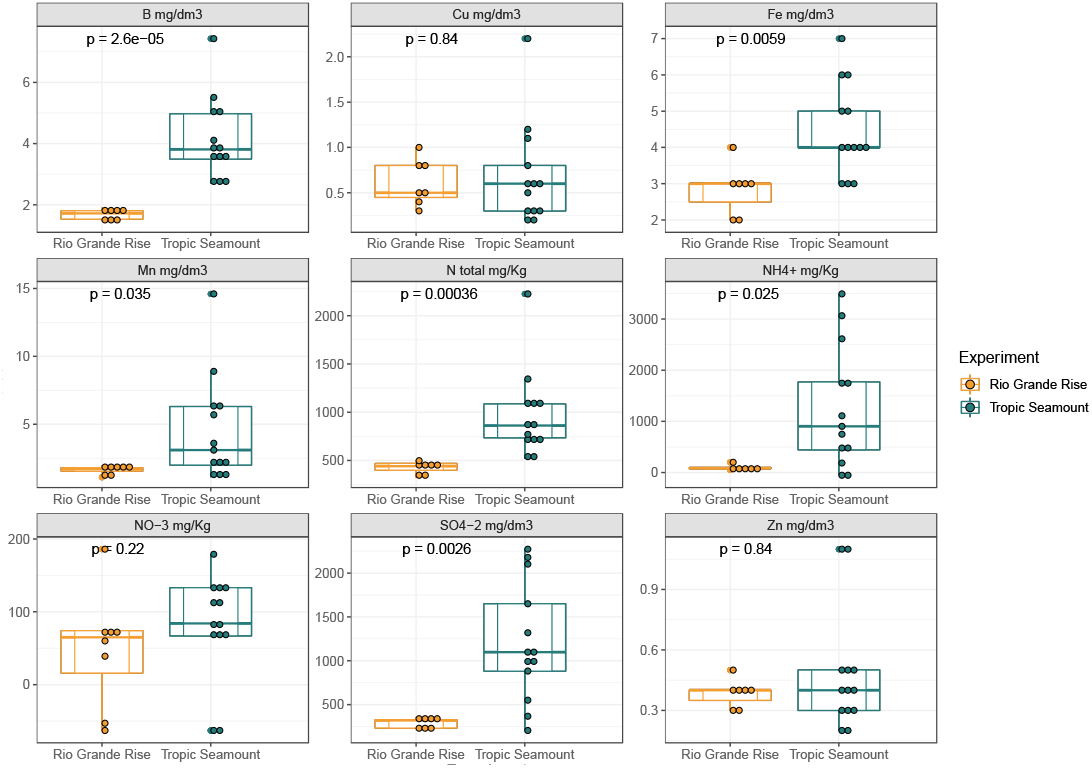
Boxplot of micronutrients (B, Cu, Fe, Mn, and Zn), nitrogen (N_total_, NH_4_, and NO_3_), and sulfur (SO_4_^−2^) compounds concentrations in sediment from Tropic Seamount (green) and Rio Grande Rise (orange). Note that the vertical scale bars differ between graphs. Pairwise comparisons were performed using the Wilcoxon test.

**Supplementary Figure 3.**
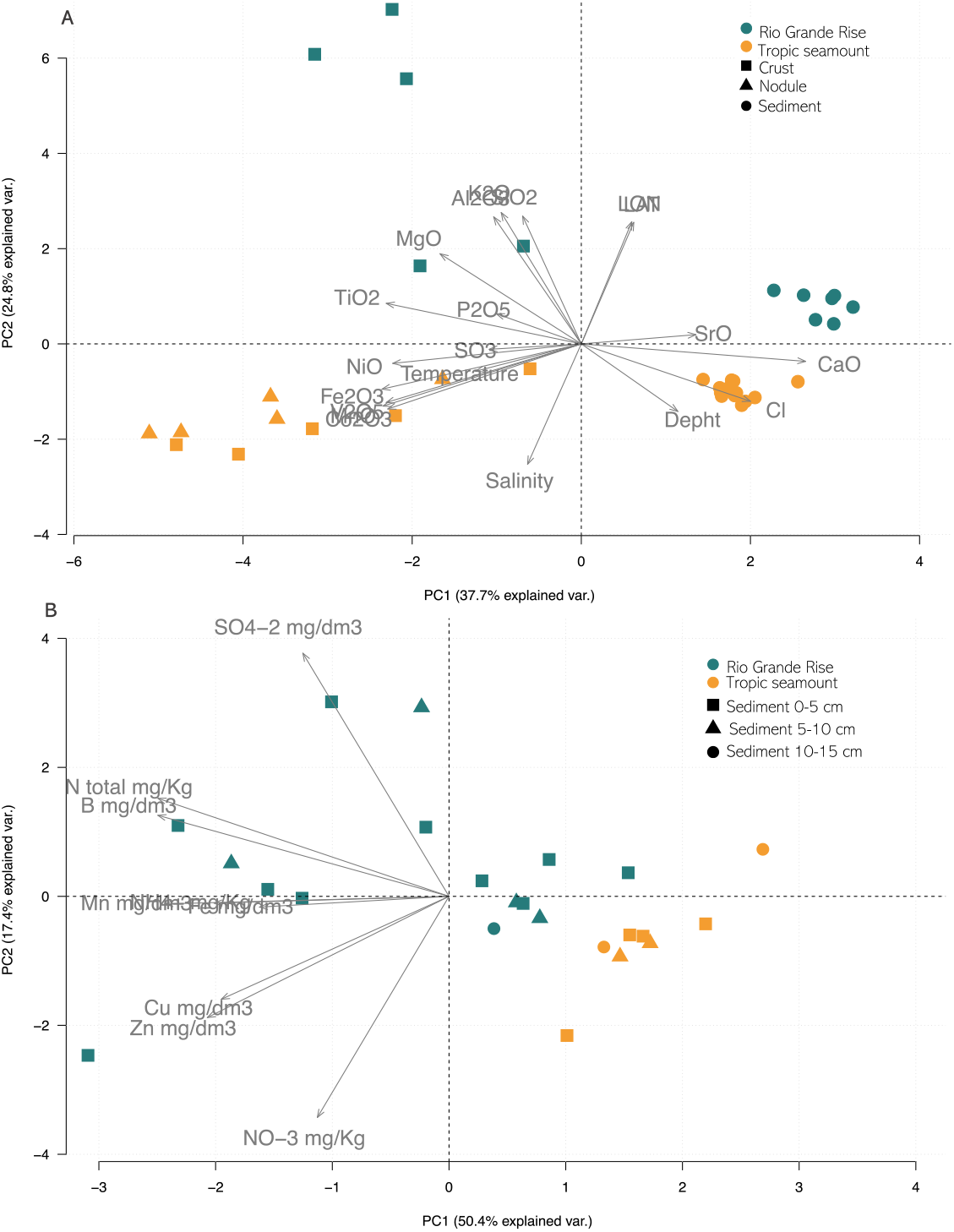
Principal component analysis samples for samples collected from Rio Grande Rise (cyan) and Tropic Seamount (orange) in the Atlantic Ocean. (a) Fe-Mn crusts (squares), nodules (triangle), and associated sediment (circle) geochemical data and oceanographic parameters are shown; (b) only sediment samples, micronutrients data are shown.

**Supplementary Figure 4.**
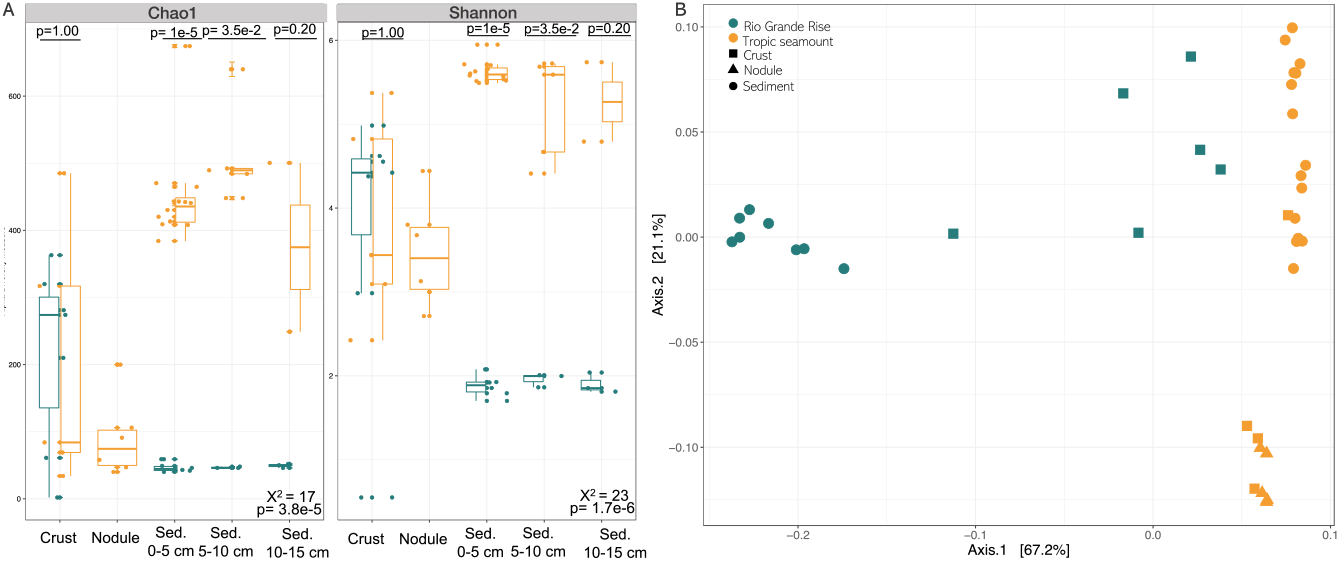
Alpha and beta diversity of microbial communities in the Fe-Mn crusts, nodules, and associated sediment from Rio Grande Rise (cyan) and Tropic seamount (orange). (a) Alpha diversity (Chao1 and Shannon indexes) medians of microbial communities. Means were compared by the Kruskal-Wallis test. Pairwise comparisons were performed using the Wilcoxon test. (b) Beta diversity and principal coordinate analysis (PCoA) are based on the ordinated weighted Unifrac normalized distance. Fe-Mn crust (square), nodule (triangle), and associated sediment (circle) from RGR and Tropic Seamount.

**Supplementary Figure 5.**
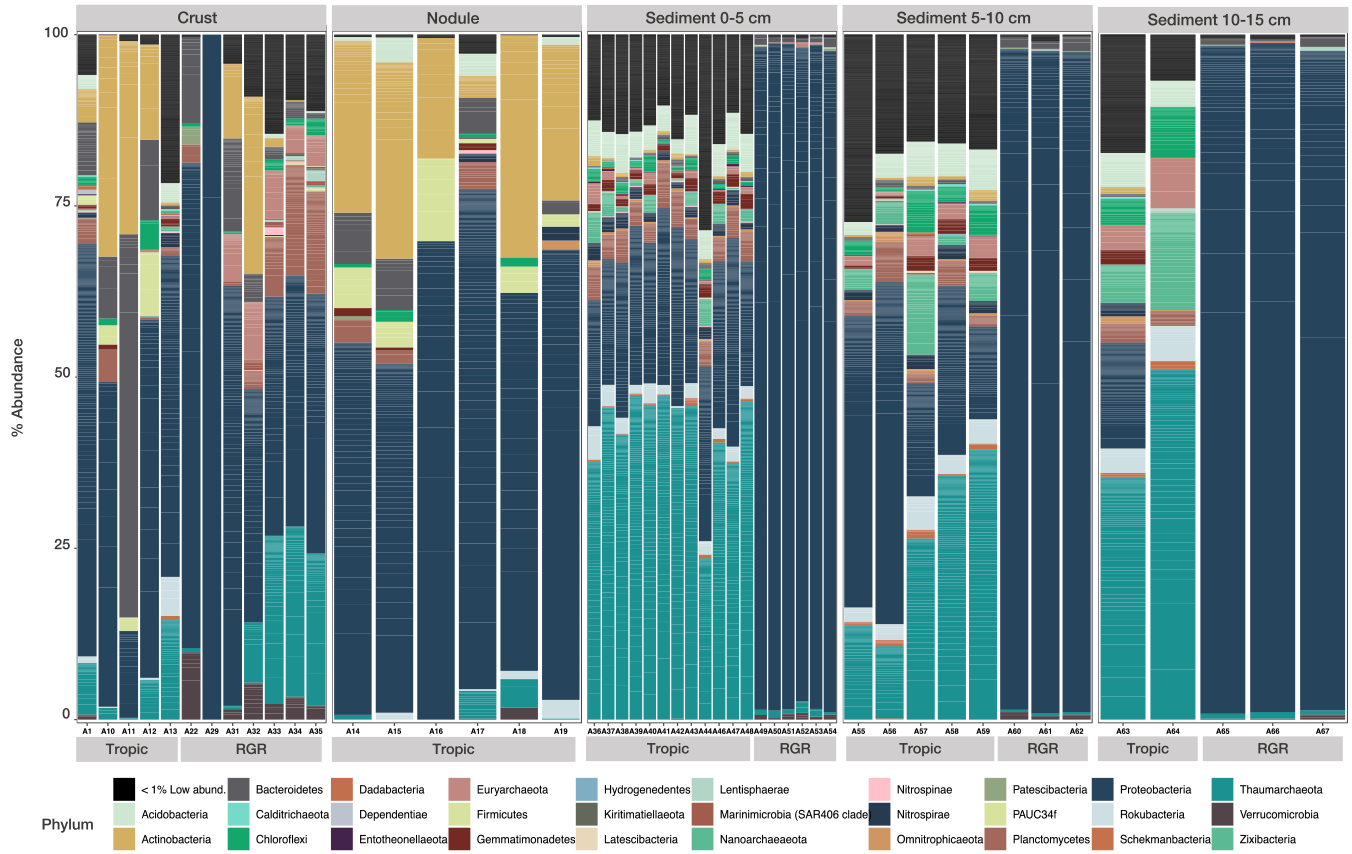
Relative abundances of bacterial and archaeal taxonomic composition for phyla in the Fe-Mn crusts, nodules, and sediment depth intervals 0-5 cm, 5-10 cm, and 10-15 cm from Tropic seamount and Rio Grande Rise. Only phyla with more than 0.1% abundance are represented. Phyla with relative abundances below 1% were grouped for the low abundance groups. Gray boxes at the top indicate sample substrates and gray boxes at the bottom differentiate samples from Tropic seamount (Tropic) and Rio Grande Rise (RGR).

**Supplementary Figure 6.**
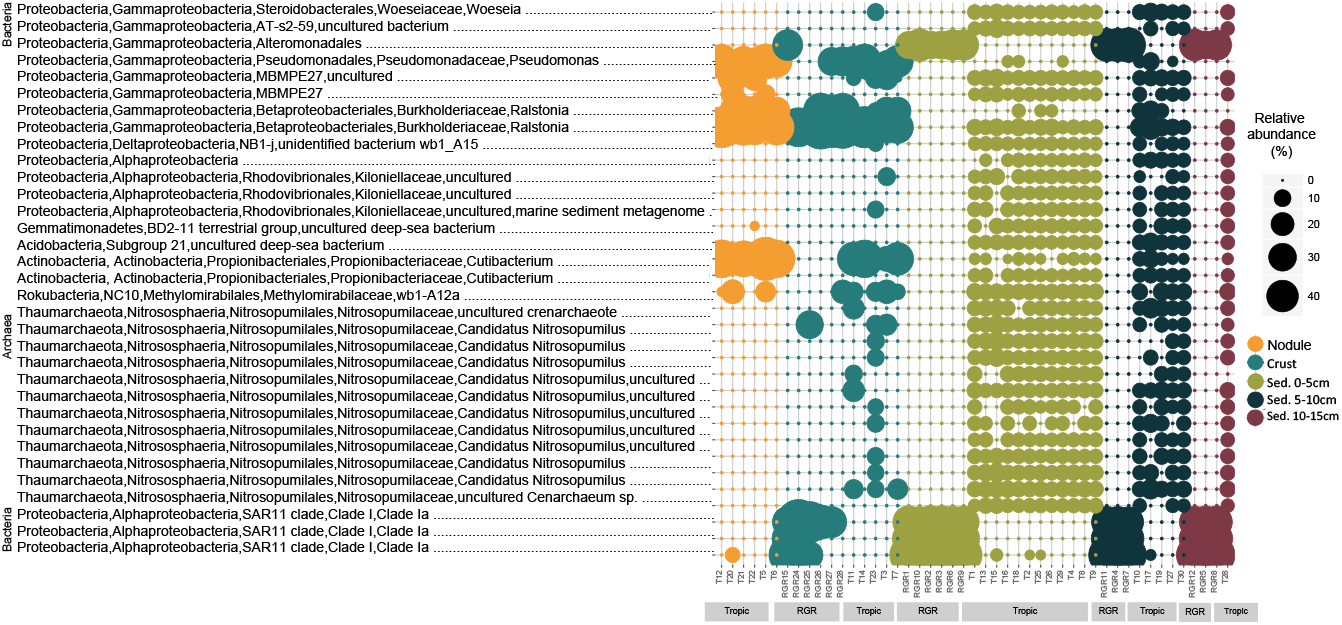
Classification of the most abundant ASVs for Archaea and Bacteria (>0.1% of relative abundance) in the Fe-Mn crusts, nodules, and sediment depth intervals 0-5 cm, 5-10 cm, and 10-15 cm from Tropic seamount and Rio Grande Rise. The size of circles is relative to the abundance of each ASV. ASVs are organized by phylum level. Gray boxes at the bottom indicate samples from Tropic seamount (Tropic) and Rio Grande Rise (RGR).

**Supplementary Table 1.** List of samples from Takuyo-Daigo seamount and Clarion-Clipperton Zone used in this study containing the sample ID, type, water depth, latitude, longitude, salinity, and temperature.

| Sample ID | Location | Sample Type | Depth (m) | LAT (°) | LON (°) | Salinity (psu) | Temperature (°C) |
| --- | --- | --- | --- | --- | --- | --- | --- |
| DRR122360 | Takuyo-Daigo Seamount | Crust | 1432 | 22.74343 | 153.26630 | 34.5 | 3.5 |
| DRR122337 | Takuyo-Daigo Seamount | Crust | 1432 | 22.7434 | 153.26630 | 34.5 | 3.5 |
| DRR122344 | Takuyo-Daigo Seamount | Crust | 1154 | 22.8517 | 156.44955 | 34.3 | 4 |
| DRR122345 | Takuyo-Daigo Seamount | Crust | 1154 | 22.8517 | 156.44955 | 34.3 | 4 |
| DRR122346 | Takuyo-Daigo Seamount | Crust | 5517 | 22.5120 | 153.21638 | 34.7 | 1.5 |
| DRR122347 | Takuyo-Daigo Seamount | Crust | 5517 | 22.5120 | 153.21638 | 34.7 | 1.5 |
| DRR122349 | Takuyo-Daigo Seamount | Crust | 2988 | 22.6809 | 153.23940 | 34.6 | 1.5 |
| DRR122350 | Takuyo-Daigo Seamount | Crust | 2987 | 22.6809 | 153.23940 | 34.6 | 1.5 |
| DRR122351 | Takuyo-Daigo Seamount | Crust | 2987 | 22.6809 | 153.23940 | 34.6 | 1.5 |
| DRR122352 | Takuyo-Daigo Seamount | Crust | 2987 | 22.6809 | 153.23940 | 34.6 | 1.5 |
| DRR122359 | Takuyo-Daigo Seamount | Crust | 5373 | 22.4865 | 153.20115 | 34.7 | 1.5 |
| DRR122341 | Takuyo-Daigo Seamount | Crust | 5373 | 22.4865 | 153.20115 | 34.7 | 1.5 |
| DRR122354 | Takuyo-Daigo Seamount | Crust | 4480 | 22.6246 | 153.21525 | 34.7 | 1.5 |
| DRR122342 | Takuyo-Daigo Seamount | Crust | 4480 | 22.6246 | 153.21525 | 34.7 | 1.5 |
| DRR122361 | Takuyo-Daigo Seamount | Sed. 0-5cm | 1432 | 22.7434 | 153.26618 | 34.5 | 3.5 |
| DRR122362 | Takuyo-Daigo Seamount | Sed. 0-5cm | 1154 | 22.8517 | 153.44955 | 34.3 | 4 |
| DRR122363 | Takuyo-Daigo Seamount | Sed. 0-5cm | 5577 | 22.5095 | 153.21815 | 34.7 | 1.5 |
| DRR122364 | Takuyo-Daigo Seamount | Sed. 0-5cm | 5577 | 22.5095 | 153.21815 | 34.7 | 1.5 |
| DRR122365 | Takuyo-Daigo Seamount | Sed. 0-5cm | 4374 | 22.6252 | 153.21815 | 34.7 | 1.5 |
| DRR122366 | Takuyo-Daigo Seamount | Sed. 0-5cm | 4374 | 22.6252 | 153.21815 | 34.7 | 1.5 |
| SRR1980774 | Clarion-Clipperton Zone | Sed. 0-5cm | 4079 | 13.8465 | 116.45983 | 34.7 | 1.5 |
| SRR1980779 | Clarion-Clipperton Zone | Sed. 0-5cm | 4078 | 13.7937 | 116.70317 | 34.7 | 1.5 |
| SRR1980781 | Clarion-Clipperton Zone | Sed. 0-5cm | 4078 | 13.7937 | 116.70317 | 34.7 | 1.5 |
| SRR1980784 | Clarion-Clipperton Zone | Sed. 0-5cm | 4084 | 13.9633 | 116.56817 | 34.7 | 1.5 |
| SRR1980789 | Clarion-Clipperton Zone | Sed. 0-5cm | 4084 | 13.9633 | 116.56817 | 34.7 | 1.5 |
| SRR1980793 | Clarion-Clipperton Zone | Sed. 0-5cm | 4054 | 13.8242 | 116.53433 | 34.7 | 1.5 |
| SRR1980797 | Clarion-Clipperton Zone | Sed. 0-5cm | 4054 | 13.8242 | 116.53433 | 34.7 | 1.5 |
| SRR1980803 | Clarion-Clipperton Zone | Sed. 0-5cm | 4076 | 13.8117 | 116.71000 | 34.7 | 1.5 |
| SRR1980805 | Clarion-Clipperton Zone | Sed. 0-5cm | 4076 | 13.8117 | 116.71000 | 34.7 | 1.5 |
| SRR1980810 | Clarion-Clipperton Zone | Sed. 0-5cm | 4111 | 13.7618 | 116.46000 | 34.7 | 1.5 |
| SRR1980814 | Clarion-Clipperton Zone | Sed. 0-5cm | 4111 | 13.7618 | 116.46000 | 34.7 | 1.5 |
| SRR1980818 | Clarion-Clipperton Zone | Sed. 0-5cm | 4111 | 13.7618 | 116.46000 | 34.7 | 1.5 |
| SRR1980823 | Clarion-Clipperton Zone | Sed. 0-5cm | 4111 | 13.7617 | 116.46033 | 34.7 | 1.5 |
| SRR1980825 | Clarion-Clipperton Zone | Sed. 0-5cm | 4111 | 13.7617 | 116.46033 | 34.7 | 1.5 |
| SRR1980824 | Clarion-Clipperton Zone | Sed. 0-5cm | 4150 | 13.8883 | 116.69000 | 34.7 | 1.5 |
| SRR1980829 | Clarion-Clipperton Zone | Sed. 0-5cm | 4150 | 13.8883 | 116.69000 | 34.7 | 1.5 |
| SRR1980841 | Clarion-Clipperton Zone | Sed. 0-5cm | 4163 | 13.9018 | 116.59000 | 34.7 | 1.5 |
| SRR1980842 | Clarion-Clipperton Zone | Sed. 0-5cm | 4163 | 13.9018 | 116.59000 | 34.7 | 1.5 |
| SRR1980843 | Clarion-Clipperton Zone | Sed. 0-5cm | 4163 | 13.9018 | 116.59000 | 34.7 | 1.5 |
| SRR1980844 | Clarion-Clipperton Zone | Sed. 0-5cm | 4166 | 13.9017 | 116.59000 | 34.7 | 1.5 |
| SRR1980848 | Clarion-Clipperton Zone | Sed. 0-5cm | 4166 | 13.9017 | 116.59000 | 34.7 | 1.5 |
| SRR1980876 | Clarion-Clipperton Zone | Nodule | 4079 | 13.8465 | 116.45983 | 34.7 | 1.5 |
| SRR1980877 | Clarion-Clipperton Zone | Nodule | 4079 | 13.8465 | 116.45983 | 34.7 | 1.5 |
| SRR1980878 | Clarion-Clipperton Zone | Nodule | 4079 | 13.8465 | 116.45983 | 34.7 | 1.5 |
| SRR1980879 | Clarion-Clipperton Zone | Nodule | 4079 | 13.8465 | 116.45983 | 34.7 | 1.5 |
| SRR1980880 | Clarion-Clipperton Zone | Nodule | 4078 | 13.7937 | 116.70317 | 34.7 | 1.5 |
| SRR1980881 | Clarion-Clipperton Zone | Nodule | 4078 | 13.7937 | 116.70317 | 34.7 | 1.5 |
| SRR1980882 | Clarion-Clipperton Zone | Nodule | 4078 | 13.7937 | 116.70317 | 34.7 | 1.5 |
| SRR1980883 | Clarion-Clipperton Zone | Nodule | 4078 | 13.7937 | 116.70317 | 34.7 | 1.5 |
| SRR1980886 | Clarion-Clipperton Zone | Nodule | 4054 | 13.8242 | 116.53433 | 34.7 | 1.5 |
| SRR1980885 | Clarion-Clipperton Zone | Nodule | 4054 | 13.8242 | 116.53433 | 34.7 | 1.5 |
| SRR1980884 | Clarion-Clipperton Zone | Nodule | 4054 | 13.8242 | 116.53433 | 34.7 | 1.5 |
| SRR1980887 | Clarion-Clipperton Zone | Nodule | 4054 | 13.8242 | 116.53433 | 34.7 | 1.5 |
| SRR1980888 | Clarion-Clipperton Zone | Nodule | 4054 | 13.8242 | 116.53433 | 34.7 | 1.5 |
| SRR1980889 | Clarion-Clipperton Zone | Nodule | 4076 | 13.8117 | 116.71000 | 34.7 | 1.5 |
| SRR1980890 | Clarion-Clipperton Zone | Nodule | 4076 | 13.8117 | 116.71000 | 34.7 | 1.5 |
| SRR1980891 | Clarion-Clipperton Zone | Nodule | 4076 | 13.8117 | 116.71000 | 34.7 | 1.5 |
| SRR1980893 | Clarion-Clipperton Zone | Nodule | 4076 | 13.8117 | 116.71000 | 34.7 | 1.5 |
| SRR1980901 | Clarion-Clipperton Zone | Nodule | 4036 | 13.7500 | 116.51333 | 34.7 | 1.5 |
| SRR1980902 | Clarion-Clipperton Zone | Nodule | 4036 | 13.7500 | 116.51333 | 34.7 | 1.5 |
| SRR1980903 | Clarion-Clipperton Zone | Nodule | 4036 | 13.7500 | 116.51333 | 34.7 | 1.5 |
| SRR1980904 | Clarion-Clipperton Zone | Nodule | 4036 | 13.7500 | 116.51333 | 34.7 | 1.5 |
| SRR1980905 | Clarion-Clipperton Zone | Nodule | 4150 | 13.8883 | 116.69000 | 34.7 | 1.5 |
| SRR1980907 | Clarion-Clipperton Zone | Nodule | 4150 | 13.8883 | 116.69000 | 34.7 | 1.5 |
| SRR1980908 | Clarion-Clipperton Zone | Nodule | 4050 | 13.8883 | 116.54883 | 34.7 | 1.5 |
| SRR1980909 | Clarion-Clipperton Zone | Nodule | 4163 | 13.9018 | 116.59000 | 34.7 | 1.5 |
| SRR1980911 | Clarion-Clipperton Zone | Nodule | 4163 | 13.9018 | 116.59000 | 34.7 | 1.5 |
| SRR1980910 | Clarion-Clipperton Zone | Nodule | 4076 | 13.8117 | 116.71000 | 34.7 | 1.5 |
| SRR1980912 | Clarion-Clipperton Zone | Nodule | 4076 | 13.8117 | 116.71000 | 34.7 | 1.5 |
| SRR1980914 | Clarion-Clipperton Zone | Nodule | 4166 | 13.9017 | 116.59000 | 34.7 | 1.5 |
| SRR1980913 | Clarion-Clipperton Zone | Nodule | 4166 | 13.9017 | 116.59000 | 34.7 | 1.5 |
| SRR1980915 | Clarion-Clipperton Zone | Nodule | 4053 | 13.8633 | 116.54667 | 34.7 | 1.5 |
| SRR1980916 | Clarion-Clipperton Zone | Nodule | 4053 | 13.8633 | 116.54667 | 34.7 | 1.5 |
| SRR1980918 | Clarion-Clipperton Zone | Nodule | 4078 | 13.7937 | 116.70317 | 34.7 | 1.5 |
| SRR1980917 | Clarion-Clipperton Zone | Nodule | 4078 | 13.7937 | 116.70317 | 34.7 | 1.5 |
| SRR1980920 | Clarion-Clipperton Zone | Nodule | 4081 | 13.7933 | 116.61983 | 34.7 | 1.5 |
| SRR1980919 | Clarion-Clipperton Zone | Nodule | 4081 | 13.7933 | 116.61983 | 34.7 | 1.5 |

**Supplementary Table 2.** List of samples from Tropic seamount and Rio Grande Rise used in this study containing the sample ID, type, water depth, latitude, longitude, salinity, temperature, and oxygen concentration.

| Sample ID | Ship Station | Location | Sample Type | Depth (m) | LAT | LON | Salinity (psu) | Temperature (°C) | Oxygen (mg/l) |
| --- | --- | --- | --- | --- | --- | --- | --- | --- | --- |
| JC142_020_012 | 298 | Tropic Seamount | Crust | 996 | 23.8912 | 20.6992 | 35.19 | 6.93 | NA |
| JC142_094_027 | 313 | Tropic Seamount | Crust | 1350 | 23.8917 | 20.6286 | 35.23 | 5.88 | NA |
| JC142_100_008 | 314 | Tropic Seamount | Crust | 1234 | 23.8591 | 20.6861 | 35.24 | 6.46 | NA |
| JC142_100_002 | 314 | Tropic Seamount | Crust | 1400 | 23.8567 | 20.6846 | 35.21 | 5.49 | NA |
| JC142_104_008 | 316 | Tropic Seamount | Crust | 1015 | 23.8537 | 20.7248 | 35.16 | 6.82 | NA |
| JC142_020_021 | 298 | Tropic Seamount | Nodule | 996 | 23.8831 | 20.6949 | 35.2 | 7.01 | NA |
| JC142_020_022 | 298 | Tropic Seamount | Nodule | 996 | 23.8831 | 20.6951 | 35.2 | 7.01 | NA |
| JC142_080_015 | 308 | Tropic Seamount | Nodule | 1200 | 23.9005 | 20.7743 | 35.24 | 6.28 | NA |
| JC142_104_004 | 316 | Tropic Seamount | Nodule | 1015 | 23.8536 | 20.7248 | 35.16 | 6.82 | NA |
| JC142_104_005 | 321 | Tropic Seamount | Nodule | 1015 | 23.8536 | 20.7248 | 35.16 | 6.82 | NA |
| JC142_104_006 | 316 | Tropic Seamount | Nodule | 1015 | 23.8536 | 20.7248 | 35.16 | 6.82 | NA |
| JC142_020_003 | 298 | Tropic Seamount | Sed. 0-5cm | 996 | 23.8833 | 20.7055 | 35.18 | 6.97 | NA |
| JC142_020_008 | 298 | Tropic Seamount | Sed. 0-5cm | 996 | 23.8833 | 20.7062 | 35.17 | 6.9 | NA |
| JC142_020_018 | 298 | Tropic Seamount | Sed. 0-5cm | 996 | 23.8827 | 20.6958 | 35.19 | 7.01 | NA |
| JC142_094_028 | 313 | Tropic Seamount | Sed. 0-5cm | 1350 | 23.8917 | 20.6286 | 35.22 | 5.9 | NA |
| JC142_080_005 | 308 | Tropic Seamount | Sed. 0-5cm | 1200 | 23.9076 | 20.7747 | 35.24 | 6.41 | NA |
| JC142_080_021 | 308 | Tropic Seamount | Sed. 0-5cm | 1200 | 23.9002 | 20.7739 | 35.23 | 6.36 | NA |
| JC142_100_010 | 314 | Tropic Seamount | Sed. 0-5cm | 1235 | 23.8589 | 20.6861 | 35.24 | 6.47 | NA |
| JC142_100_005 | 314 | Tropic Seamount | Sed. 0-5cm | 1233 | 23.8567 | 20.6846 | 35.21 | 5.47 | NA |
| JC142_104_004 | 316 | Tropic Seamount | Sed. 0-5cm | 1015 | 23.8536 | 20.7248 | 35.16 | 6.82 | NA |
| JC142_104_009 | 316 | Tropic Seamount | Sed. 0-5cm | 1015 | 23.8537 | 20.7248 | 35.16 | 6.81 | NA |
| JC142_104_014 | 316 | Tropic Seamount | Sed. 0-5cm | 1013 | 23.8536 | 20.7267 | 35.16 | 6.82 | NA |
| JC142_116_014 | 321 | Tropic Seamount | Sed. 0-5cm | 1018 | 23.8581 | 20.7419 | 35.16 | 6.82 | NA |
| JC142_080_005 | 308 | Tropic Seamount | Sed. 5-10cm | 1200 | 23.9076 | 20.7747 | 35.24 | 6.41 | NA |
| JC142_100_005 | 314 | Tropic Seamount | Sed. 5-10cm | 1233 | 23.8567 | 20.6846 | 35.21 | 5.47 | NA |
| JC142_104_004 | 316 | Tropic Seamount | Sed. 5-10cm | 1015 | 23.8536 | 20.7248 | 35.16 | 6.82 | NA |
| JC142_104_014 | 316 | Tropic Seamount | Sed. 5-10cm | 1013 | 23.8536 | 20.7267 | 35.16 | 6.82 | NA |
| JC142_116_014 | 321 | Tropic Seamount | Sed. 5-10cm | 1018 | 23.8581 | 20.7419 | 35.16 | 6.82 | NA |
| JC142_104_014 | 316 | Tropic Seamount | Sed. 10-15cm | 1013 | 23.8536 | 20.7267 | 35.16 | 6.82 | NA |
| JC142_116_014 | 321 | Tropic Seamount | Sed. 10-15cm | 1018 | 23.8581 | 20.7419 | 35.16 | 6.82 | NA |
| RGR1_DR10_01 | S0480 | Rio Grande Rise | Crust | 702 | 30.86793 | 35.9999 | 34.41 | 6.61 | 7.09 |
| RGR1_DR11 | S0484 | Rio Grande Rise | Crust | 775 | 31.0094 | 36.1787 | 34.44 | 6.76 | 6.99 |
| RGR1_DR11_02 | S0484 | Rio Grande Rise | Crust | 775 | 31.0094 | 36.1787 | 34.44 | 6.76 | 6.99 |
| RGR1_DR15 | S0509 | Rio Grande Rise | Crust | 638 | 30.63737 | 35.7391 | 34.46 | 6.97 | 6.94 |
| RGR1_DR17_P | S0510 | Rio Grande Rise | Crust | 686 | 30.70133 | 35.7184 | 34.46 | 6.97 | 6.94 |
| RGR1_DR17_G | S0510 | Rio Grande Rise | Crust | 686 | 30.70133 | 35.7184 | 34.46 | 6.97 | 6.94 |
| RGR1_BC1_I | S0450 | Rio Grande Rise | Sed. 0-5cm | 963 | 31.41668 | 35.8441 | 34.28 | 4.53 | 7.12 |
| RGR1_BC2_I | S0451 | Rio Grande Rise | Sed. 0-5cm | 1155 | 32.41668 | 34.8854 | 34.28 | 4.53 | 7.12 |
| RGR1_BC3_I | S0468 | Rio Grande Rise | Sed. 0-5cm | 1552 | 30.87007 | 35.7169 | 34.52 | 3.06 | 5.92 |
| RGR1_BC3_II | S0468 | Rio Grande Rise | Sed. 0-5cm | 1230 | 30.8698 | 35.9547 | 34.35 | 3.51 | 6.45 |
| RGR1_BC3_III | S0468 | Rio Grande Rise | Sed. 0-5cm | 685 | 30.93683 | 36.2621 | 34.64 | 2.96 | 6.13 |
| RGR1_BC4_I | S0474 | Rio Grande Rise | Sed. 0-5cm | 1566 | 30.83867 | 35.8113 | 34.38 | 6.05 | 7.09 |
| RGR1_BC4_II | S0474 | Rio Grande Rise | Sed. 5-10cm | 1552 | 30.87007 | 35.7169 | 34.52 | 3.06 | 5.92 |
| RGR1_BC4_III | S0474 | Rio Grande Rise | Sed. 5-10cm | 1230 | 30.8698 | 35.9547 | 34.35 | 3.51 | 6.45 |
| RGR1_BC6_I | S0500 | Rio Grande Rise | Sed. 5-10cm | 1566 | 30.83867 | 35.8113 | 34.38 | 6.05 | 7.09 |
| RGR1_BC7_I | S0504 | Rio Grande Rise | Sed. 10-15cm | 1552 | 30.87007 | 35.7169 | 34.52 | 3.06 | 5.92 |
| RGR1_BC7_II | S0504 | Rio Grande Rise | Sed. 10-15cm | 1230 | 30.8698 | 35.9547 | 34.35 | 3.51 | 6.45 |
| RGR1_BC7_III | S0504 | Rio Grande Rise | Sed. 10-15cm | 1566 | 30.83867 | 35.8113 | 34.38 | 6.05 | 7.09 |

**Supplementary Table 3.** Sequences were denoised using DADA2 (Callahan et al., 2016) with the parameters listed below.

| **Fe-Mn deposit** | **Ocean** | **p-trim-left-f** | **p-trim-left-r** | **p-trunc-len-f** | **p-trunc-len-r** |
| --- | --- | --- | --- | --- | --- |
| Tropic Seamount | Atlantic | 280 | 297 | 19 | 18 |
| Rio Grande Rise | Atlantic | 270 | 280 | 19 | 18 |
| Clarion-Clipperton Zone | Pacific | 250 | 250 | - | - |
| Takuyo-Daigo Seamount | Pacific | 0 | 250 | - | - |

**Supplementary Table 4.** Results of principal component analysis using environmental variables (longitude, latitude, depth, salinity, and temperature) samples from Tropic Seamount, Rio Grande Rise, Takuyo-Daigo Seamount, and Clarion-Clipperton Zone. Principal component loadings > 0.70 are shown in bold.

| Variable | PC1 | PC2 | PC3 |
| --- | --- | --- | --- |
| Longitude | **0.93** | 0.14 | 0.21 |
| Latitude | **-0.79** | 0.52 | 0.3 |
| Depth | **0.94** | -0.18 | 0.13 |
| Salinity | -0.45 | -0.87 | 0.18 |
| Temperature | **-0.97** | -0.06 | 0 |
| % Explanation | 70.4 | 21.7 | 0.3 |

**Supplementary Table 5.**
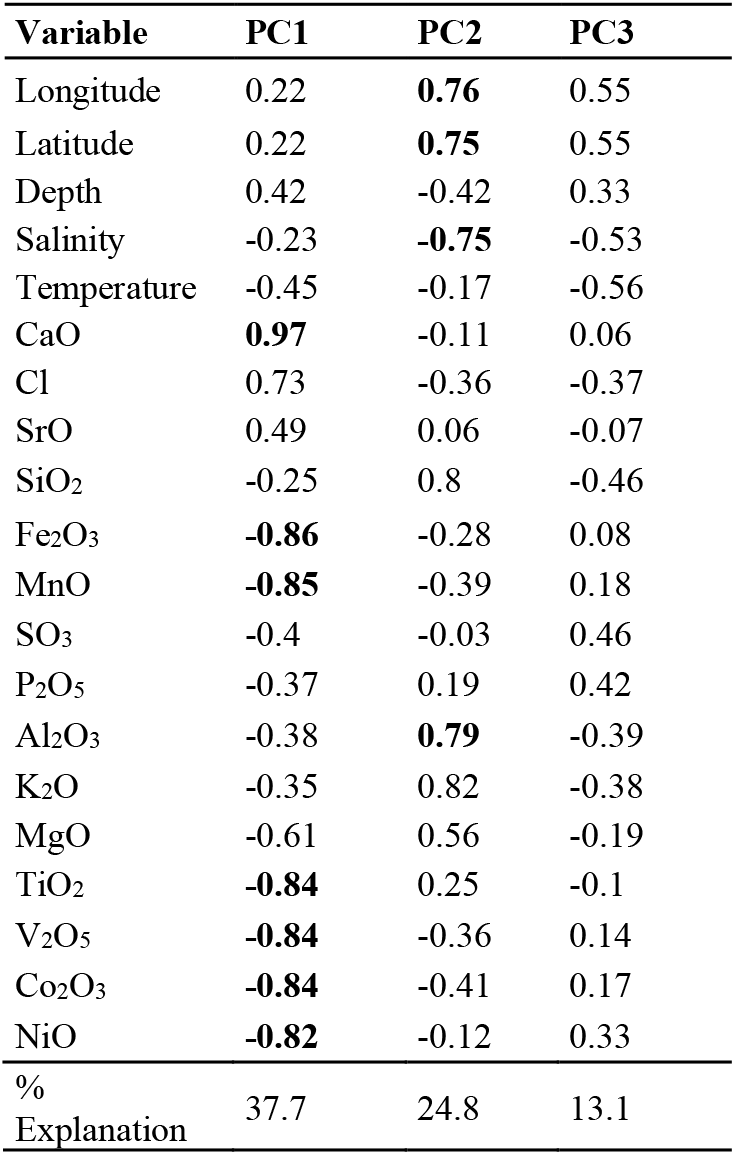
Principal component analysis using environmental variables and chemical composition of Fe-Mn crusts, nodules, and sediments (longitude, latitude, depth salinity, CaO, Cl, SrO, SiO_2_, Fe_2_O_3_, MnO, SO_3_, P_2_O_5_, Al_2_O_3_, K_2_O, MgO, TiO_2_, V_2_O_5_, Co_2_O_3_, and NiO) from Tropic Seamount and Rio Grande Rise. Principal component loadings >0.70 are shown in bold.

**Supplementary Table 6.**
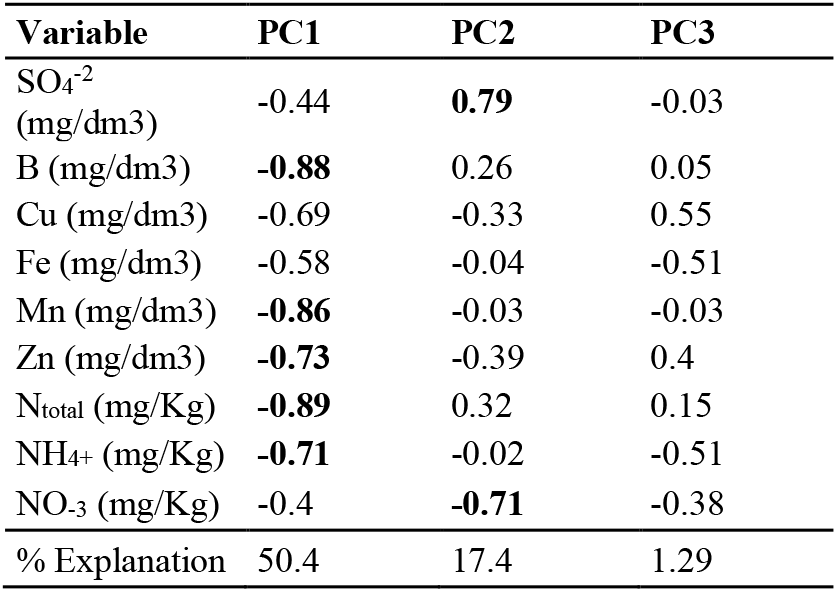
Principal component analysis of nitrogen and sulfur compounds (N_total_, NH_4+_, NO_-3_, and SO_4_^−2^) and micronutrients (B, Cu, Fe, Mn, and Zn) from Tropic Seamount and Rio Grande Rise. Principal component loadings > 0.70 are shown in bold.

**Supplementary Table 7.** Sum, mean, standard error (S. error) for number of ASVs, Chao1 and Shannon indexes for Fe-Mn crust, nodule, and sediment samples from Atlantic and Pacific Oceans.

| Ocean | Sample type | Chao1 |  |  | Shannon |  |  |
| --- | --- | --- | --- | --- | --- | --- | --- |
|  |  | Sum | Mean | S. error | Sum | Mean | S. error |
| Atlantic | Crust | 3218.25 | 268.19 | 52.26 | 51.67 | 4.31 | 0.32 |
|  | Nodule | 542.00 | 90.30 | 24.31 | 20.76 | 3.4 | 0.26 |
|  | Sed. 0-5 cm | 6037.78 | 317.78 | 46.05 | 83.65 | 5.52 | 0.41 |
| Pacific | Crust | 2051.00 | 146.5 | 11.28 | 66.9 | 4.78 | 0.09 |
|  | Nodule | 33123.00 | 920.08 | 35.19 | 211.34 | 5.91 | 0.04 |
|  | Sed. 0-5 cm | 25299 | 937.00 | 106.49 | 158.60 | 5.88 | 0.11 |

**Supplementary Table 8.**
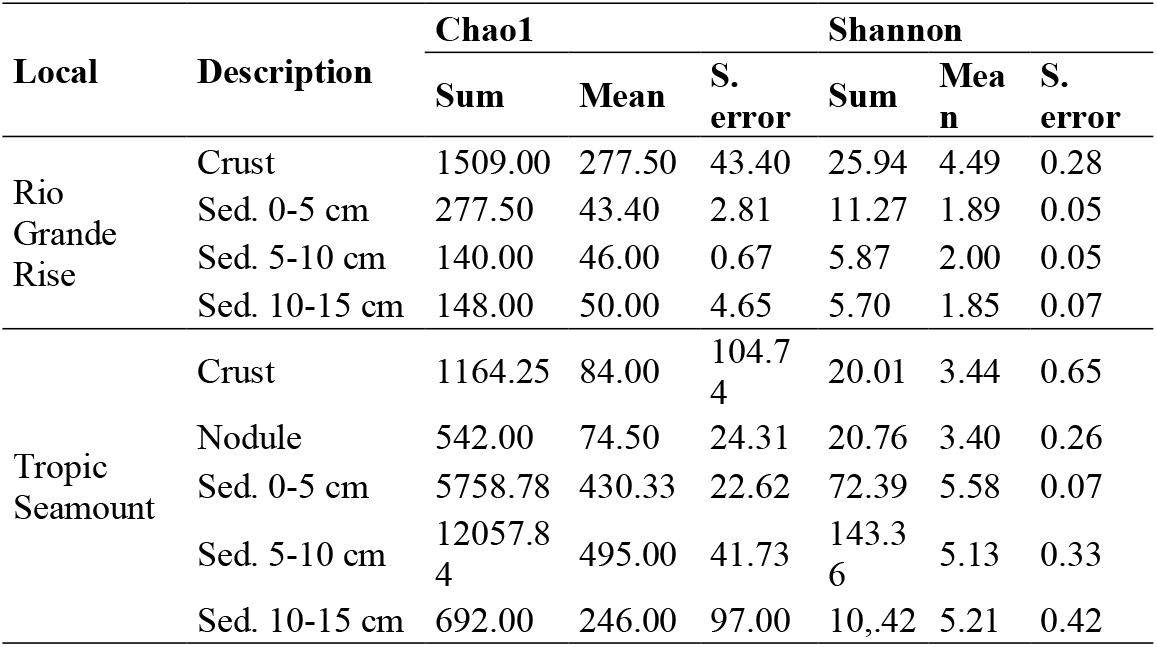
Sum, mean, Standard error (S. error) for number of ASVs, Chao1, and Shannon indexes for Fe-Mn crust, nodule, and sediment samples from Tropic Seamount and Rio Grande Rise.

